# Identification of Myeloid Protein Kinase C – Epsilon as a Novel Atheroprotective Gene

**DOI:** 10.1101/2024.12.09.627650

**Authors:** Alexis T. Wells, Ramon Bossardi Ramos, Michelle M. Shen, Redwan H. Binrouf, Anna E. D’Amico, Michelle R. Lennartz

## Abstract

**BACKGROUND:** Atherosclerosis is a chronic inflammatory disease driven by macrophages (MØ). Protein Kinase C – epsilon (PKCɛ) is a serine/threonine kinase involved in diverse cellular processes including migration, growth, differentiation, and survival. PKCɛ acts in a context dependent manner within the heart, however, its role in atherosclerosis is unknown.

**METHODS:** Bone marrow derived MØ from global PKCɛ knockout mice were tested for lipid retention and cytokine secretion. Public geneset analysis assessed raw counts of PRKCE in human atheromas to determine translational relevance. A LysM Cre PKCɛ^fl/fl^ (mɛKO) mouse was developed to study the impact of myeloid PKCɛ on atherosclerosis. After confirming myeloid selective PKCɛ deletion, human-like hypercholesterolemia was induced and multiple metrics of atherosclerosis were compared in WT and mɛKO plaques. RNA sequencing was used to provide unbiased insight into possible mechanisms by which PKCɛ regulates atherosclerosis.

**RESULTS:** Public geneset analysis of human atherosclerotic plaque tissue revealed that PKCɛ expression is inversely correlated with plaque vulnerability. Similarly, peritoneal MØ from WT hypercholesterolemic mice have significantly lower PKCɛ expression, providing a translational rational for generation of the mɛKO mouse. qPCR revealed no differences between genotypes in the expression of genes related to atherosclerosis, at either steady state or upon lipid loading, suggesting that loss of PKCɛ does not fundamentally change the basal state and that differences seen are a result of a more complex pathway. Comparing descending aorta and aortic root plaques from WT and mɛKO hypercholesterolemic mice revealed that mɛKO plaques are larger, have larger foam cells and regions of necrosis, and thinner collagen caps. Upon lipid loading in vitro and in vivo, mɛKO MØ retained significantly more cholesterol and lipid droplets than WT; Gene Ontology suggests higher expression of genes related to endocytosis in mɛKO MØ compared to WT.

**CONCLUSIONS:** PKCɛ expression is decreased in vulnerable human plaques and decreases in mouse MØ upon lipid loading. mɛKO plaques are larger and exhibit markers of vulnerability. With no differences in scavenger receptor (SR) expression, the impact of PKCɛ deletion is more subtle than simple SR dysregulation. RNA sequencing implicates higher expression of genes involved in endocytosis and mɛKO MØ have significantly more lipid-containing endosomes. The data define the atherophenotype of mɛKO mice and demonstrate that PKCɛ restricts lipid uptake into MØ by a mechanism independent of SR expression. Taken together, these studies identify PKCɛ as a novel atheroprotective gene, laying the foundation for mechanistic studies on the endocytic signaling networks responsible for the phenotype.

**GRAPHIC ABSTRACT:** 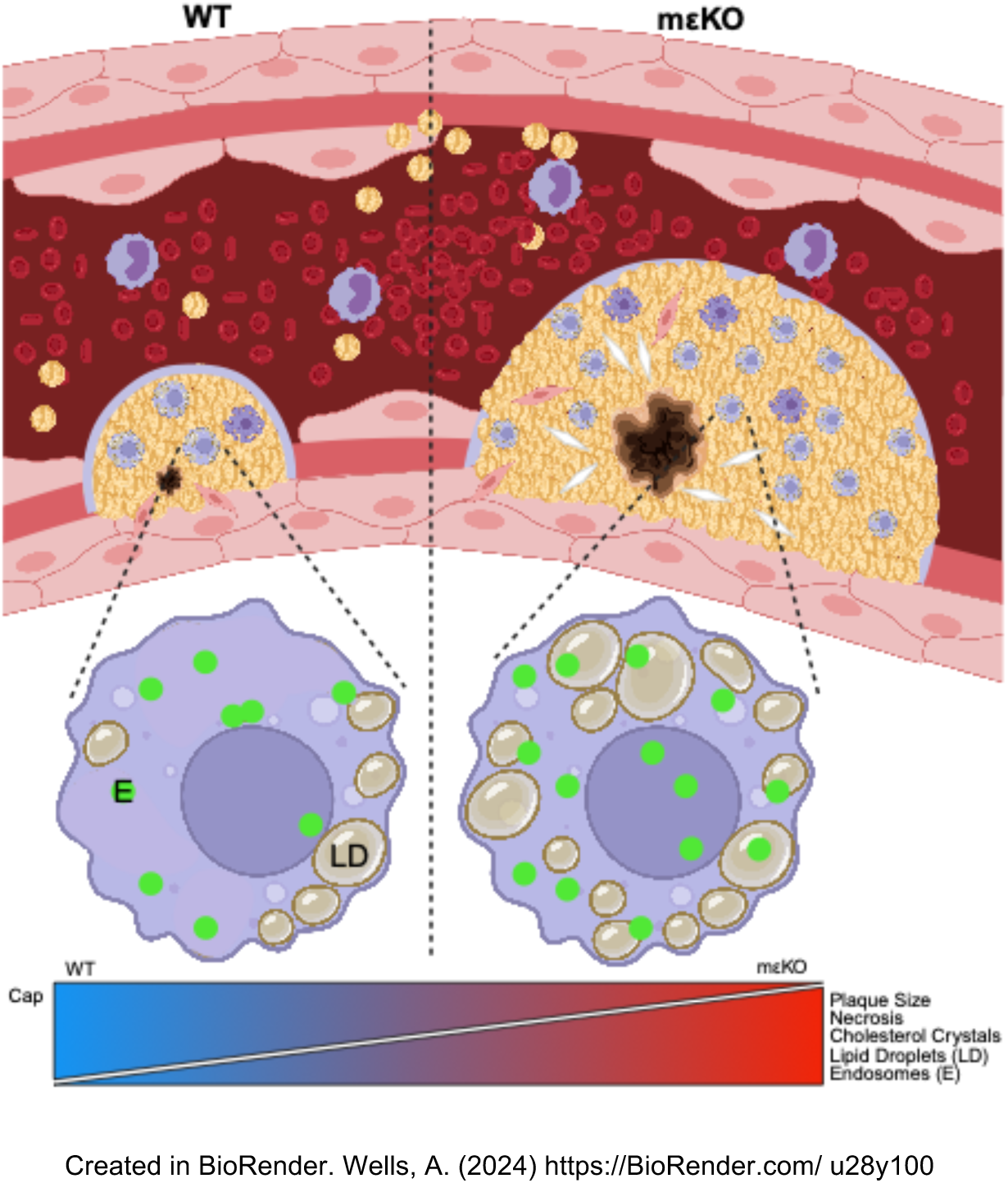

**Highlights:**

- A novel murine model in which PKCε is floxed (PKCε^fl/fl^) on both alleles haas been generated, backcrossed, and deposited into Jackson Laboratories (B6(Cg)-*Prkce^tm1.1Akv^*/LennMmjax). Then, PKCε^fl/fl^ LysM Cre (mεKO) mice have been generated.
- Deletion of PKCε has no basal affects on other PKC isoforms, lipid handling markers, or inflammatory markers.
- Upon (in vitro) lipid loading of BMDM or isolating lipid loaded pMACs from hypercholesterolemic mice (in vivo), mεKO BMDMs retain more cholesterol.
- Long term (16 wk) hypercholesterolemis generates larger lesions, more necrosis, and thinner plaque caps (i.e., a more vulnerable plaque phenotype) in mεKO mice.
- RNA sequencing suggests PKCε regulates endocytosis mechanisms post initial uptake of LDL and has been confirmed in a model of synchronized endocytosis.
- These findings identify myeloid PKCε as a novel atheroprotective molecule whose expression impacts multiple metrics of atherosclerosis. This opens new avenues of investigation to uncover the PKCε signaling network responsible for the phenotype. Understanding how myeloid PKCε slows atherosclerosis development will aid in the development of novel preventative and therapeutic treatments.

## Introduction

Cardiovascular disease (CVD) is the leading cause of death worldwide, with atherosclerosis being a major contributor.^1,2^ Despite the widespread use of statins, CVD remains a major health care problem, accounting for 1 in 5 deaths in the U.S. (2022) and 12% of health care costs ($378 billion in 2017-2018).^3,4^ New strategies and targeted therapies are needed to combat these statistics, reducing atherosclerosis and the catastrophic effects (heart attack and stroke) of plaque rupture.

There is a cascade of events that occur during the development of atherosclerosis (Figure 2A).^5,6^ Macrophages (MØ) migrate to sites of endothelial damage, extravasate into the intima, and internalize modified and non-modified low-density lipoprotein (LDL) through scavenger receptors or LDLR. At later stages, cells within the plaque become apoptotic and, if not removed by efferocytosis, contribute to the necrosis prevalent in advanced plaques.^7^ While plaque size may vary, the more critical factor in assessing vulnerability is plaque composition.^5,6,8^ That is, human vulnerable plaques have thin fibrous caps, embedded collagen covered by new plaque growth, are foam cell rich, and have large necrotic cores.^6,8^ This constellation of factors produces chronic inflammation that can result in thrombosis, leading to angina, myocardial infarction (MI), stroke, and even death.^5,9,10^

Previously, we analyzed gene expression in human carotid and femoral plaques and found that proximal regions of carotid plaques had increased expression of MØ activation markers, including FcγR, SR-A1, TNFα, and PKCδ.^11^ Protein Kinase Cs are a family of master kinases, linked to virtually every known signaling network. Our subsequent work identified protein kinase C epsilon (PKCɛ), a PKCδ related family member, as an inflammatory signaling node.^12^ Its involvement in IgG-mediated phagocytosis, respiratory burst, and NF-κB activation is well documented.^4,13–21^ This suggests that PKCɛ can moderate excessive inflammation but *how* it accomplishes this is a fundamental unanswered question. As atherosclerosis is a chronic inflammatory disease, we investigated the role of macrophage PKCε in plaque development.

PKCs act in protection and pathology, depending on context.^17,22–25^ PKCɛ has been linked to inflammation in patients with atherosclerosis, hypertension-induced heart failure, and conversely, to protection against ischemia-reperfusion injury and myocardial infarction.^22,23,25,26^ In vascular smooth muscle cells, PKCɛ inhibits apoptosis, increases MMP-2 and -9 expression, and induces migration. ^27^ In cardiomyocytes, PKCɛ regulates the interaction between conexin43 and a potassium channel subunit (Kir6.1) within the mitochondria, preventing hypoxia-induced cell death.^28^ In MØ, PKCɛ has been linked to phagocytosis of aggregated LDL and resistin signaling via TLR4/MyD88, thus increasing lipid accumulation and inflammation.^23,27,29,30^ In contrast, a large randomized clinical trial using high-dose adenosine (activator of PKCɛ) during acute MI found a significant *reduction* in infarct size.^27,31^ Thus, the role of PKCɛ in cardiovascular disease is still an open question.

PKCɛ is perhaps best known as an oncogene, overexpressed in many cancers where it enhances tumor cell migration, invasion, and resistance to apoptosis.^32–37^ If applied to MØ and atherosclerosis, these properties could be atheroprotective. That is, PKCɛ may support MØ recruitment into early plaques, regulate the uptake/retention/ efflux of cholesterol, promote cell survival extending MØ lifetime for increased efferocytosis, and protect against metrics of atherogenesis that contribute to MI and heart failure. Thus, the same PKCɛ processes that are pathological in cancer may slow the development of atherosclerosis making it a potential therapeutic target. as a therapeutic agent for several diseases. Its expression at steady state may sustain homeostasis but its decrease or overexpression produces pathology. Yet, the role of myeloid PKCɛ in atherosclerosis is unknown.

In vivo studies indicate that loading MØ with LDL decreases PKCɛ expression, which parallels RNAseq data from human plaques (PKCε is significantly lower in advanced plaques). This would suggest that PKCε levels decrease as atherosclerosis progresses and implicates PKCε in lipid handling. This hypothesis is supported by studies with MØ from PKCε KO mice, which revealed that PKCε null MØ retain more lipid and secrete more TNFα, both pro-inflammatory responses. To study the impact of MØ PKCɛ in atherosclerosis, we generated PKCɛ^fl/fl^ mice and crossed them to LysM Cre, generating a myeloid specific PKCε knockout mouse (mɛKO). Using the AAV8-PCSK9 model to induce hypercholesterolemia in mɛKO and their siblings lacking LysM Cre (i.e., “wild type” with respect to PKCɛ expression), we applied a variety of human plaque metrics. Plaques from mɛKO mice were more advanced with evidence of vulnerability. RNA sequencing of MØ from hypercholesterolemic WT and mɛKO mice suggests that PKCε regulates signaling pathways related to endocytosis, that may restrict lipid retention, slowing disease progression. Taken together, this work presents multiple lines of evidence implicating MØ PKCɛ in atheroprotection and implicates endocytosis signaling as a potential mechanism.

## MATERIALS and METHODS

### Sex as a Biological Variable/Limitations of the Study

Atherosclerosis disproportionately affects human males vs pre-menopausal women.^38^ As 8-24 weeks in mice correspond to ~15-25 years in humans,^39^ and mice undergo changes similar to menopause at 9-12 months,^40^ female mice are still cycling at 8-24 weeks. Thus, to maximize the chances of seeing phenotypic differences, this study was limited to males. It should be noted that while males and females were included in the first cohort, the females did not remain hypercholesterolemic for the 16 weeks and did not produce plaques. The response of female mice to AAV8-PCSK9 strategy is reported to be lower than that of males, so the virus will have to be titrated^41^ to ensure long-term hypercholesterolemia before determining the effect of MØ PKCɛ on plaque progression in females.

### Reagents

ACK lysis buffer: 0.5 M NH_4_Cl, 1 mM KHCO_3_, and 0.1 mM EDTA (pH 7.4). Bone marrow media (BMM): DMEM containing 10% FBS, 20% L Cell media, 1% Na Pyruvate, 1% Glutamax, 0.03% sodium bicarbonate, 1% nonessential amino acids, and 50 μg/ml gentamicin. Cell media: BMM lacking L cell media. Sucrose lysis buffer/P^3^I: 50 mM Tris-HCl, pH 7.4, 0.25 M sucrose, 2.5 mM DTT, 2.5 mM EDTA, containing P^3^I [5 mM benzamidine, 50 mg/mL leupeptin, 50 mg/mL aprotinin, 50 mg/mL trypsin inhibitor, 5 mg/mL pepstatin, 1 mM PMSF, 20 mM NaF, 1 mM Na_3_VO_4_, 1 mM para-nitro-phenylphosphate, 5 mM imidazole (Sigma)]. TBST: 5 mM Tris, pH 7.4 containing 150 mM NaCl and 0.1% Tween® 20. Flow buffer: PBS, 2% FBS, 2mM EDTA. Flow blocking buffer: flow buffer containing 10% rat serum and 5 µg/ml 2.4G2 FcR blocking Ab).

### Publicly Available RNA-Seq Data (GSE1104140)

The RNA-seq data was retrieved from GSE104140. Raw counts of PRKCE, as a measure of unmodified gene expression, in diffuse intimal thickening (defined by AHA as normal intima^42^; and advanced calcified or non-calcified fibroatheroma. Data was analyzed using iDEP custom R code for differential expression (limma-voom). Principle component analysis (PCA) emphasizes the variation between groups as well as the similarities within each group.

### Insoluble Immune Complexes

10 µl of 10 mg/mL BSA (Sigma A0281) was added to 300 µl of 2 mg/mL rabbit anti-BSA IgG (Sigma, B1520) in Optimem^TM^ buffer (ThermoFisher Scientific, 31985070). Complexes formed by rotation (1 hour, 37°C). Complexes were pelleted and washed twice with Optimem, and resuspended to the initial volume. 15 µl of this stock solution was added/1 x 10^6^ BMDM.

### Mice

All animal procedures were approved by the Albany Medical Center Institutional Animal Care and Use Committee protocol #1903003 and #2204004 (breeding) and #2304003 (AAV8 and high fat diet). Cells from both males and females were used for *in vitro* studies; there were no effects of sex or age (3-6 months) on experimental outcome. Cells obtained from one mouse represent one independent experiment. All animals/cells were identified by ear-tag (not genotype) to minimize unconscious bias. The lab manager identified animals to be used in each experiment. Genotypes were revealed after all data was collected.

### Global PKCε Knockout Mice (ɛKO)

PKCε^+/−^ heterozygotes on the C57BL/6 background were purchased from Jackson Laboratory (*B6.129S4-Prkce^tm1Msg^/J* stock# 004189; Bar Harbor, ME) and bred in the Albany Medical Center Animal Resources Facility. Mice were fed a standard chow, housed under regular 12/12 light-dark cycles at a temperature of 20-26°C, and a humidity level of 20-70%. Global deletion of PKCɛ in mice leads to noticeably smaller offspring with an immunodeficiency causing ɛKO mice to succumb to infections and fall out of lineage. Heterozygotes were crossed to produce the wild-type (C57BL/6) and PKCε^−/−^ (ɛKO) siblings.

### Myeloid Selective PKCε Knockout Mouse (mɛKO)

PKCɛ^fl/fl^ mice were generated by IVF at Taconic Labs (Germantown, NY) from sperm provided by The Genome Editing and Animal Models Core at the University of Wisconsin (Madison) Biotechnology Center. This C57BL/6 congenic strain was deposited at The Jackson labs (B6(Cg)-*Prkce^tm1.1Akv^*/LennMmjax stock# 69556). PKCɛ^fl/fl^ females were then crossed to LysM Cre males (Jackson Laboratory, Strain #004781 B6.129P2-*Lyz2^tm1(cre)Ifo^*/J) to produce LysM Cre^+/−^ PKCɛ^fl/fl^ pups. To maintain the colony PKCɛ^fl/fl^ females were crossed to LysM Cre^+/−^ PKCɛ^fl/fl^ males to produce PKCɛ^fl/fl^ offspring, some of which lacked LysM Cre (designated WT) and siblings expressing hemizygous LysM Cre (**<u>m</u>**yeloid selective PKCɛ **<u>k</u>**nock**<u>o</u>**ut, designated mɛKO). A subset of this colony carried the ZsGreen Cre reporter gene (B6.Cg-*Gt(ROSA)26Sor^tm6(CAG-ZsGreen1)Hze^*/J maintained as heterozygous) and was used to confirm specificity of Cre expressing MØ in mɛKO plaques [as well as in blood (neutrophils and MØ) and splenocytes (B and T cells). Zs green carrying animals are indicated by green symbols on graphs; expression of ZsGreen had no impact on plaque metrics. Mice were fed a standard chow, housed under regular 12/12 light-dark cycles at a temperature of 20-26°C, and a humidity level of 20-70%. Compared to the parental C57BL/6 strain, there are no obvious size, behavioral, or health issues with either the WT or mɛKO animals or in animals carrying/expressing the ZsGreen gene.

### Generation of Hypercholesterolemic Mice

Cohort animals were identified and their eartags recorded. Weights of 8-week-old mice were recorded at the time of retro-orbital injections of AAV8-PCSK9 (1×10^12^ gc/200 µl sterile PBS). Animals were given a high fat (HFD, “Western”) diet (Inotiv, TD.88137) for 4 or 16 weeks. At two weeks post-injection/HFD, and at the time of sacrifice, blood was collected (submandibular bleed or portal vein pull) for plasma cholesterol measurements (Fujifilm Healthcare Americas Corp Cholesterol E kit, NC9138103); ≥500 mg/dL was considered hypercholesterolemic. mɛKO mice are indistinguishable from their WT counterparts. Breeding LysMCre hemizygote studs to LysMCre negative dams produced litters of 7-13 animals that were approximately 50% LysMCre^+/−^ making it relatively easy to generate sufficient numbers of each genotype for the atherosclerosis cohorts.

The final experimental conditions were chosen from the results of a pilot experiment. Pilot experiments: 16 mice (8 male, 8 female; two timepoints 9- and 16-weeks post AAV8/HFD). Females had high plasma cholesterol at 2 weeks that decreased over time. Males remained hypercholesterolemic for the duration. At 9 weeks, no animals had visible plaque. At 16 weeks, males, but not females, had developed plaques. The lack of plaques in females correlates with their lack of sustained hypercholesterolemia due to the AAV8-PCSK9 model dosage. Thus, subsequent cohorts were male only, harvesting cells 4 weeks and tissue 16 weeks after AAV8/HFD.

### Elicited Peritoneal MØ (pMACs)

Mice were injected i.p. with 3 mL sterile 3% Thioglycolate solution. Animals were euthanized after four days and pMACs recovered by peritoneal lavage with 10mL PBS and red blood cells removed via ACK lysis.

### Bone Marrow-derived MØ (BMDM)

Differentiated macrophages were derived from bone marrow stem cells as previously described.^12^ Briefly, mice were euthanized via isoflurane and cervical dislocation. Femurs and pelvises were removed and flushed for bone marrow collection. Red blood cells were lysed with ACK lysis buffer and the recovered stem cells were differentiated in BMM. BMDM were used 7-9 days after harvesting.

### ELISA assays

C57BL/6 and ɛKO BMDM (1.5 × 10^6^/well, 6 well plate) were incubated with or without immune complexes for 2 hours. Culture media was collected and IL6 and TNFα was quantified using BioLegend ELISA Max^TM^ kits (431301 and 430901) per the manufacturer’s instructions. Results are reported as as pg/mL net release.

### Western Blotting

<u>Brains</u> were homogenized (sucrose lysis buffer/P^3^I) using a KINEMATICA Polytron PT 10 20 3500 115v. The homogenate was sonicated on ice 2 × 10 s (Sonics & Materials Vibra-cell VC High Intensity Ultrasonic Processor) and centrifuged (13,000 x g, 10 minutes, 4°C) to remove particulates. 3 µg of protein (Bradford Assay) was used for western blots. <u>Spleens</u> were minced in PBS and passed through a 70 µm cell strainer (CELLTREAT, 229483). Erythrocytes were ACK lysed and splenocytes recovered by centrifugation (400 x g, 10 minutes, 4°C). Splenocytes were solubilized in 400 µl sucrose lysis buffer, followed by sonication on ice (2 × 10 s). After clarification by centrifugation (13,000 x g, 10 minutes, 4°C), protein was quantified (Bradford Assay); 100 µg was loaded/lane. <u>BMDMs and pMACs</u> were suspended in 100 µl Sucrose Lysis Buffer/P^3^I, and sonicated for 7 s on ice. Samples were clarified by centrifugation (13,000 x g, 10 min 4°C), and protein was quantitated; 300 µg of protein/lane was used for western blots.

Proteins were separated on a 10% PAGE under reducing conditions. The Bio-Rad Precision Plus Protein Kaleidoscope ladder (1610375) served as the molecular weight standards. Gels were transferred onto a PVDF membrane using an iBlot® system. Membranes were blocked with 5% milk in 1X TBST (1 hour) and washed 3 times with TBST. Antibodies were diluted in 3% BSA/TBST (Major Resources Table) and allowed to bind overnight (4°C with rocking); GAPDH served as the loading control. The following day, membranes were washed 5 times with TBST, probed with an HRP secondary antibody (1:10,000 in 3% BSA/TBST, Major Resources Table) for 1 hour RT while rocking. Membranes were washed (5 times with TBST) and developed using Bio-Rad’s Clarity Western ECL Substrate kit (1705060) according to the manufacturer’s instructions. Membranes were imaged using a Bio-Rad ChemiDoc^TM^ MP Imaging System. Densitometry was used to quantify genotype differences in PKC expression. TIFF files were opened in FIJI and the Analyze → Gels function was used to generate peaks to the corresponding bands. Peaks were separated and the wand tool used to quantify the area of each band. For each band the background was subtracted, normalized to GAPDH and their WT counterparts. Fold change relative to C57BL/6 or WT was reported.

### Blood Collection for Complete Blood Count (CBC)

Blood was collected from mice via the portal vein into 1mL syringes containing 100 µl heparin. CBC analysis was performed using a Heska ElementHT5 Veterinary Hematology Analyzer (Heska Corporation, Loveland, Colorado). The percent composition of total white blood cells was calculated and reported. The total number of circulating RBC, WBC, lymphocytes, monocytes, neutrophils, eosinophils, and basoplils were collected and reported as Cells (10^6^/μl).

### qPCR

RNA from 1×10^6^ MØ were extracted with PureLink™ RNA Mini Kit (ThermoFisher Scientific, 12183018A); RNA purity and concentration was determined by Nanodrop. RNA was reversed transcribed using qScript cDNA SuperMix (QuantaBio, 95048-500). cDNA was then used for qPCR. Each 20 µl reaction contained 10 µl Quanta PerfeCTa Syber Green Fast Mix (QuantaBio, 95071-012), 8 nMol of forward and reverse primers (Origene), and 50 ng cDNA. Thermocycler Ct values were exported into an Excel file and the relative expression (2^−ΔCt^) was calculated and reported. All samples were normalized to β-actin (see Major Resources Table for primer sequences).

### Flow Cytometry

50 - 100 µl of BMDM or pMAC (at 1 × 10^7^/mL) were blocked (30-60 min, ice) before antibody staining (45 min, ice in the dark; Major Resources Table). After staining, cells were resuspended in 500 µl of flow buffer and read on a Bio-Rad ZE5 Cell Analyzer driven by Everest software. Data was analyzed with FlowJo 10.10.

### In Vitro Lipid Loading

*Human highly-oxidized-LDL* (Kalen Biomedical, #770252): 5×10^5^ BMDM were incubated with 50 µg/mL ox-LDL (24 hours, complete media), washed with PBS, and processed for flow cytometry, cholesterol retention, or western blotting.

*Ac-LDL* (ThermoFisher Scientific, #L23380): 5.0 × 10^5^ BMDM were incubated with 5 µg/mL A488 Acetylated LDL (30, 60 minutes, complete media), washed with PBS, resuspended in flow buffer for flow cytometery.

### Quantitation of Lipid Droplets in BMDM from Global PKCε Knockout (εKO) Mice

Initial studies (Figure 1) were done using BMDM from the global PKCε knockout mice (εKO). 5.0 × 10^4^ C57BL/6 or ɛKO BMDM were plated on coverslips in 24 well plates. Acetylated LDL (Kalen BioMedical, 770201-6; 360 µg/mL in BMM) was added. After 4 hours, the cells were washed with PBS and fixed (4% PFA, 15 min, RT). Coverslips were washed with PBS, stained with BODIPY (ThermoFisher Scientific, D3922, 30 min) and mounted. Z stacks were collected, 3-D reconstitutions were made, and droplet number and area was quantified using Imaris® software.

**Figure 1:**
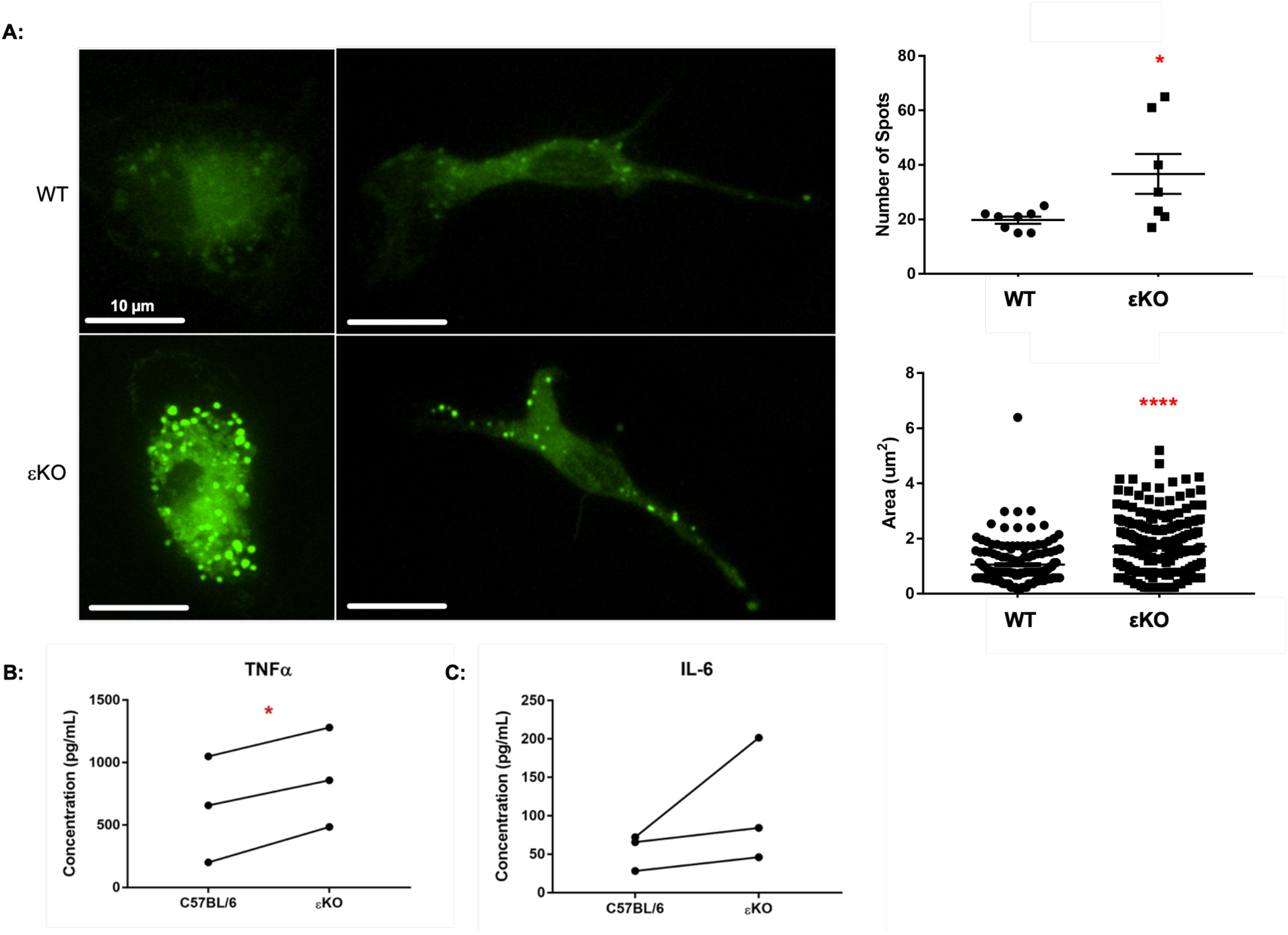
MØ from global PKCε knockout (εKO) mice retain more/larger lipid droplets and release more TNFα. (**A**) C57BL/6 and εKO BMDM were treated with AcLDL (360 µM) for 4h, fixed and lipid droplets stained with BODIPY. Z-stacks were collected, 3-D reconstructions made, and number of lipid droplets and droplet area was quantified (each symbol represents one cell/droplet n = 3 independent experiments). Data are presented as ± SEM. (**B,C**) C57BL/6 and εKO BMDM were incubated with or without immune complexes (2 h). Media was collected and TNFα and IL-6 was quantified by ELISA. Net TNFα and IL-6 release is represented (n = 3). Statistical analysis was preformed by two-tailed unpaired t-test (**A**) and two-tailed paired t-test in (**B,C**). Scale bars: 10 µm. *\*p <* 0.05; *\*\*\*\*p <* 0.0001.

### Quantitation of Cholesterol Retention

BMDM were lipid loaded (24 h, ox-LDL), washed and held at −80°C until analyzed. Total and free cholesterol was quantified from the cell lysates using the Cholesterol Ester-Glow^TM^ kit (Promega, J3190) according to the manufacturer’s instructions. Data was normalized to the total amount of protein collected from cell lysates. Technical replicates were averaged, and cholesterol ester content calculated as the difference between total and free cholesterol.

### Lipid Droplet Quantitation in Foam Cells

Four weeks after induction of hypercholesterolemia, elicited pMACs were harvested, resuspended in Dulbecci’s modified Eagle’s medium (DMEM) and adhered in 8-well chamber slides (Falcon, 354108) (30 minutes). Lipid droplets were visualized with LipidTOX (Invitrogen, H34476; 1:100) (30 minutes, RT) and mounted using ProLong Glass Antifade Mountant with NucBlue (Invitrogen, P36981). Merged images were created using FIJI software.

### Tissue Collection and Processing

Studies were terminated by isoflurane euthanasia and cervical dislocation. Blood was collected from the portal vein for plasma cholesterol determination. Animals were exsanguinated with 10 mL warm PBS through the left ventricle using a syringe and 25-gauge needle at constant pressure. Hearts and aortic arches/descending aortas were collected.

<u>Descending aortas</u> were collected from the base of the ascending aorta to the common iliac artery, keeping the branches of the brachiocephalic, left common carotid, and left subclavian arteries intact. Aortas were fixed (4% PFA overnight, RT) and stored at 4°C in PBS/Thimerosal. *En face* preparations were made by opening the lumen, pinning the tissue flat, and staining with 0.3% ORO (15 minutes, RT). After staining, arches were washed with 70% EtOH for 1 minute followed by 2 washes of 1X PBS for 5 minutes. Tissue area was defined by the aortic arch and 3 mm of the descending aorta (no plaque was present beyond 3 mm). Plaque area was quantitated in FIJI using the color thresholding macro and normalized to tissue area. The percent lesion coverage was reported.

<u>For aortic roots</u>, hearts were cut parallel to the tip of the atria. The upper half, containing the aortic root was placed in a cryomold containing optimal cutting temperature compound (OCT) and held at −80°C until sectioned. A total of 25 charged slides with 3-4 10 μm thick sections per slide were collected from each mouse as previously described.^43^ Briefly, sequential sections were put on individual slides, such that sections 1 to 25 went on slides 1 to 25. Sections 26 to 50 were placed on slides 1 to 25, respectively, and so on. This generated 25 slides, each containing multiple sections separated by ~250 μm and spanning the entire plaque length. Six slides (e.g. Slides 1, 5, 9, etc.) were chosen for H & E staining. Contiguous slides (e.g. 2,6,10, etc.) were stained with ORO. Sirius red staining was done on serial slides (e.g. 3,7,11, etc.) and the next slides in the series (e.g. 4,8,12, etc.) for F4/80. This strategy produced multiple datapoints (6 slides x multiple measurements per slide) for each plaque feature, with contiguous slides allowing for visualization of different features in close proximity.

### H&E Staining and Analysis

To quantify lesion size and necrosis, 6 slides (see above) were chosen, with the first beginning at the base of the aortic root (the first site of plaque deposition). Fresh tissue sections were fixed for 3 minutes in a solution comprised of 100% EtOH, 40% PFA, 100% Glacial acetic acid, and diluted with diH_2_O to make a 4% PFA solution. Fixed sections were dehydrated in 70% EtOH followed by 95% EtOH (1 minute), washed in running tap H_2_O and diH_2_O (30 s), stained with Hematoxylin (4 minutes), and washed with tap H_2_O until clear. Stained samples were then clarified for 15 seconds, washed with tap H_2_O, placed in bluing agent for 1 minute and then washed with tap H_2_O. After Hematoxylin staining, tissues were placed in 95% EtOH followed by Eosin-Y stain for 15 s each. To wash away excess Eosin samples were placed in multiple changes of 95% EtOH (x2) and 100% EtOH (x3) for 1 minute each. Tissues were cleared in xylene, mounted in Cytoseal XYL, and left to dry overnight prior to imaging. Images were analyzed with NDP.view 2 software. The lesion and necrotic regions were outlined, and the areas were recorded. Areas were then averaged between sections of the same mouse and the average lesion area per mouse was reported. Percent necrosis was then calculated by dividing the necrotic area by lesion area per mouse.

### ORO Staining and Analysis

Sections consecutive to those stained with H&E were taken to quantify average foam cells size. Slides were fixed (4% PFA, 10 minutes), washed, and dried with 60% isopropyl alcohol. Sections were stained with 0.3% ORO (20 minutes), rinsed with H_2_O, counter-stained with Hematoxylin (1 minute) and mounted in Gelvatol. Slides were imaged within 24 hours of staining. Images were analyzed with NDP.view 2 software. Regions of foam cells were measured for area and divided by the number of nuclei within that area (number of nuclei obtained from H&E sections). This was performed at 3-5 regions on all sections, averaged for each mouse and reported as average foam cell size/animal.

### Sirius Rred Staining and Analysis

Consecutive sections to those stained with ORO were taken to quantify cap thickness and percent collagen. Slides were fixed in 70% EtOH for 3 minutes, washed with diH_2_O and stained in 0.1% Sirius red/Direct 80 saturated in an aqueous solution of picric acid (1 hour, room temperature). Tissues were washed (2 x 0.5% acetic acid, 5 minutes), dehydrated in 100% EtOH and cleared with xylene. Slides were mounted with Cytoseal XYL and dried overnight prior to imaging. Images were analyzed with FIJI software. The percent collagen content was calculated by normalizing the collagen area to the lesion area and was reported per mouse. The plaque cap thickness was taken from 3-5 regions along each plaque and averaged per section. If no cap was present, then the thickness was recorded as 0. Average cap thickness was then divided by the lesion area and was reported per mouse. The thickest and thinnest cap regions were then extracted and assessed for genotype differences.

### RNA Ssequencing (GSE291828)

8 week-old WT and mɛKO mice were administered with AAV8-PCSK9 and given a chow or Western diet (as previously described) for 4 weeks and elicited pMACs were harvested. RNA was isolated from each mouse using TRIzol (Invitrogen, 15596026) according to manufacturer instructions. Sequencing was done as previously described, but with the mm10 gemone.^44^ Briefly, all RNA samples passed quality control (>8.6 RIN) and the RNA library was performed using the polyA selection method. RNA samples were sequenced by GENEWIZ using the illumina HiSeq system in a 2 x 150-bp configuration. Files were trimmed using Trim Galore (GitHub repository, https://github.com/FelixKrueger/TrimGalore) and aligned/annotated with Rsubread’s v1.5.3 to the mm10 genome. Genes above 0.5 counts-per-million in at least 3 samples were kept in the analysis. Differential expression analysis was performed using limma-voom and significance was defined as *P* value < 0.05. Principal component analysis (PCA) indicates a distinct gene expression profile between conditions using R. Functional enrichment analysis was performed using WebGestalt (https://www.webgestalt.org/), and Metascape (version 3.5). Notably, between WT and mɛKO atheroprone mice (AAV8-PCSK9 + Western diet/HFD), Gene Ontology revealed the *Import Into Cell* GO family of differentially regulated genes was within the top 20 GO families. This family includes genes related to phagocytosis, endocytosis, and receptor-mediated endocytosis. From which, a heatmap was generated using raw counts in R.

### Imaging

<u>Lipid loaded C57BL/6 and ɛKO BMDM</u>: Cells were imaged on an Olympus IX81-DSU spinning-disc confocal microscope at 60X oil with a Hamamatsu electron multiplying charge-coupled device camera, driven by MetaMorph software (Molecular Devices, Sunnyside, CA). Raw images were then processed using Imaris Software. <u>Aortic</u> <u>arches</u>. Olympus SZ61 Dissecting Microscope equipped with an HD camera & monitor system from Excelis HDS was used to acquire images of aortic arches. Raw images were processed with FIJI software. <u>Aortic root:</u> H&E, ORO, and Sirius red images were aquired on a high-Speed, High-resolution Digital Slide Scanner. Hamamatsu NanoZoomer 2.0RS. Images were collected with a 40X air objective. F4/80 images were aquired on an Olympus BX51 microscope (Olympus, Tokyo, Japan) equipped with a QImaging Retiga 2000R digital camera (QImaging, Surrey, British Columbia, Canada) and CellSens software (Olympus Life Science, Waltham, MA). Images were collected with a 40X air objective. Raw images were processed with NDP.view 2 or FIJI software.

*In vivo lipid loaded WT and mɛKO pMAC*. Images were aquired on an Olympus BX51 microscope (Olympus, Tokyo, Japan) equipped with a QImaging Retiga 2000R digital camera (QImaging, Surrey, British Columbia, Canada) and CellSens software (Olympus Life Science, Waltham, MA). Images were collected with a 40X air objective. Raw images were processed with FIJI software.

### Statistics

All experiments were conducted in a blinded fashion using ear-tags for identification. Once all data was analyzed, the code was broken to run statistics. Data are expressed as mean ± SEM of independent experiments or animals; *n* is noted in the figure legends. Limma-voon was used to assess differential expression. Comparison of means between groups was evaluated by one-way ANOVA or two-way ANOVA with Tukey multiple comparisons, 2-tailed paired or unpaired students t-test, or Mann-Whitney U test. Bland-Altman test was used for interobserver variability. A p-value less than 0.05 was considered statistically significant. All statistical analyses were performed using GraphPad Prism, SPSS, and R. Representative images were chosen based on those closest to the mean.

### Study Approval

All animals received humane care in accordance with the NIH Guide for the Care and Use of Laboratory Animals. All procedures were approved by the Albany Medical Center Institutional Animal Care and Use Committee protocol #1903003 (εKO breeding), #2204004 (mεKO breeding), and #2304003 (WT and mεKO AAV8 and high fat diet).

### Data Availability

The public geneset data used to assess PRKCE read counts in early and advanced stages of atherosclerosis can be found on NCBI under GSE104140. RNA sequencing performed for this manuscript can be found on NCBI under GSE291828. Please see the Major Resources Table in the Supplemental Materials.

## RESULTS

### ɛKO BMDM and a human atheroma RNAseq dataset suggest PKCɛ may be atheroprotective, regulating lipid dynamics

Based on our work demonstrating the role of PKCɛ in vesicular trafficking and FcR-mediated phagocytosis,^12^ we hypothesized that PKCɛ is involved in the uptake of modified lipids. To test this, we incubated bone marrow-derived macrophages (BMDM) from wild type (C57BL/6) and global PKCɛ knockout (ɛKO) mice with acetylated LDL (acLDL) and quantified lipid droplet number and area. ɛKO BMDM contained significantly more, and larger, lipid droplets, implicating PKCɛ in lipid handling (Figure 1A). Secondly, we know that the vulnerable regions of human carotid plaques have elevated expression of Fc receptors (FcR) and TNFα, that PKCɛ is activated upon FcR ligation, and that MØ treated with oxLDL-immune complexes become foam cells.^11,45^ The impact of PKCɛ expression on cytokine release is unknown. Thus, we treated C57BL/6 and ɛKO BMDM with immune complexes (to mimic the uptake of immune complexes present at high levels in humans with atherosclerosis) and quantified release of TNFα and IL6, two cytokines linked to atherosclerosis.^46,47^ Surprisingly, we found that secretion of TNFα, but not IL-6, was significantly *higher* in ɛKO cells (Figure 1B,C). This suggests that PKCɛ restricts TNFα, but not IL-6 release. That MØ lacking PKCɛ retain more lipid and release more TNFα suggests that PKCɛ limits foam cell formation and restricts the release of inflammatory cytokines, properties that could be protective, effectively slowing atherosclerosis progression.

To assess the translational potential of our (mouse) results, we searched for human plaque datasets but found none reporting protein expression. However, PRKCE (PKCɛ gene) raw counts were extracted from GSE104140, a curated RNAseq dataset obtained from carotid endarterectomy tissue that includes compares diffuse intimal thickening, (“stage 0”, a preclinical control^48,49^, n=9) with two types of late stage plaques, calcified (n=9) and soft atheromas (n=14). Raw counts were used because they provide the most basic and unbiased representation of gene expression. The principle component analysis (PCA) demonstrates the differential expression patterns of atheromas between each group as well as the similarities within each group (Figure 2B). Using R, raw counts of PRKCE per person were plotted and assessed for statistical differences. Compared to diffuse intimal thickening, advanced plaques have significantly lower PRKCE counts. While there was no statistical difference between the calcified and soft atheromas, counts trend lower in the advanced soft plaque, suggesting that PKCɛ expression is inversely correlated with plaque stage (Figure 2B).^50^

**Figure 2:**
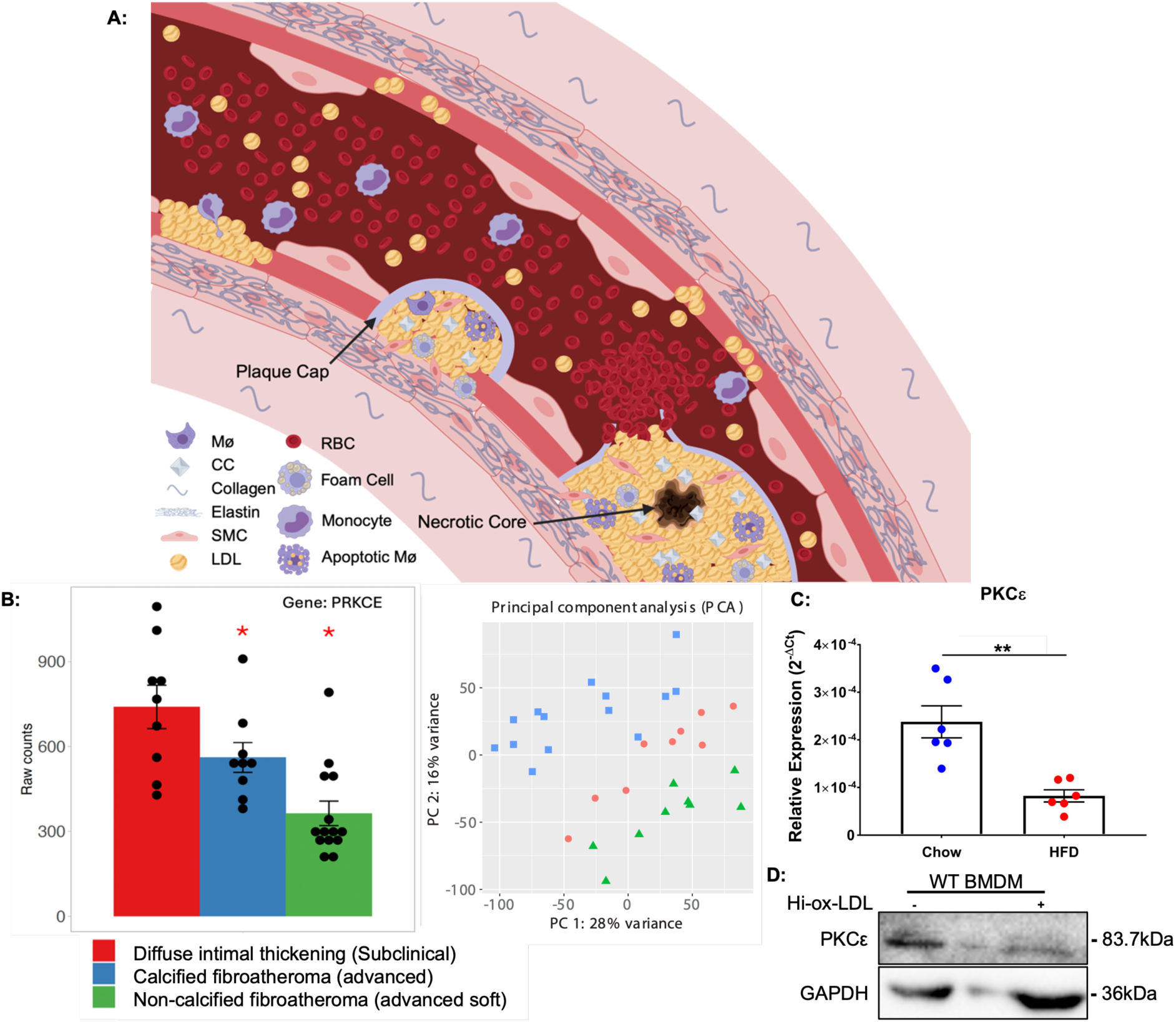
PKCε expression is significantly lower in advanced human plaques and is recapitulated in murine pMACs from hypercholesterolemic mice. (**A**) Schematic of atherogenesis. Created in BioRender. Wells, A. (2024) https://BioRender.com/b96z554. (**B**) RNAseq of human atherosclerotic plaques (GSE104140) plotting PRKCE raw counts (left) and the corresponding principal component analysis (PCA) (right) (n = 9-14 samples). (**C**) qPCR of PKCε in elicited pMACs from chow and HFD fed WT mice administered with AAV8- PCSK9 for4-wks (n=3). (**D**) Western blot of BMDM treated with Hi-ox-LDL (50 µg/mL, 24h). Data are presented as ± SEM. Statistical analysis was performed by Limma-voon differential expression analysis (**B**) and two-tailed unpaired t-test (**C**). *\*\*p <* 0.01.

Given that PKCε null MØ retain more lipid, and advanced human plaques have lower PKCε expression, we asked if MØ from hypercholesterolemic mice have lower PKCɛ. The AAV8-PCSK9 plus Western diet strategy was used for inducing hypercholesterolemia^51^; control animals received AAV8-PCSK9 but were fed chow. Elicited pMACs were harvested 4 weeks after AAV8 injection and PKCε expression quantified by PCR. Compared to chow fed, PKCε expression was significantly lower in pMACs from HFD mice (Figure 2C). To test the hypothesis that lipid loading decreases PKCε protein,we quantified PKCε expression in control MØ (no lipid) and foam cells produced upon incubation with highly oxidized low-density lipoprotein (Hi-ox-LDL); lipid loading decreased PKCε protein levels by ~ 73 ± 2% (n=2, Figure 2D). Thus, looking at mouse and human, mRNA and protein, we see a consistent decrease in PKCε in advanced plaques and hypercholesterolemic MØ, suggesting that the presence of PKCε in MØ limits lipid retention and, possibly, development of atherosclerosis. Testing this required a mouse in which PKCε is selectively deleted from myeloid cells, an mεKO mouse.

### LysM Cre deletes PKCɛ from Macrophages

We generated a PKCɛ^fl/fl^ mouse^52^ and crossed it to LysM Cre. LysM Cre deletes flox’d genes in the myeloid lineage, primarily macrophages and neutrophils, with little to no effect on the lymphoid lineage.^53^ PKCɛ^fl/fl^ dams were mated with LysM Cre^+/−^ PKCɛ^fl/fl^ studs to produce PKCɛ^fl/fl^ siblings that were LysM Cre negative or hemizygous (Figure 3A).

**Figure 3:**
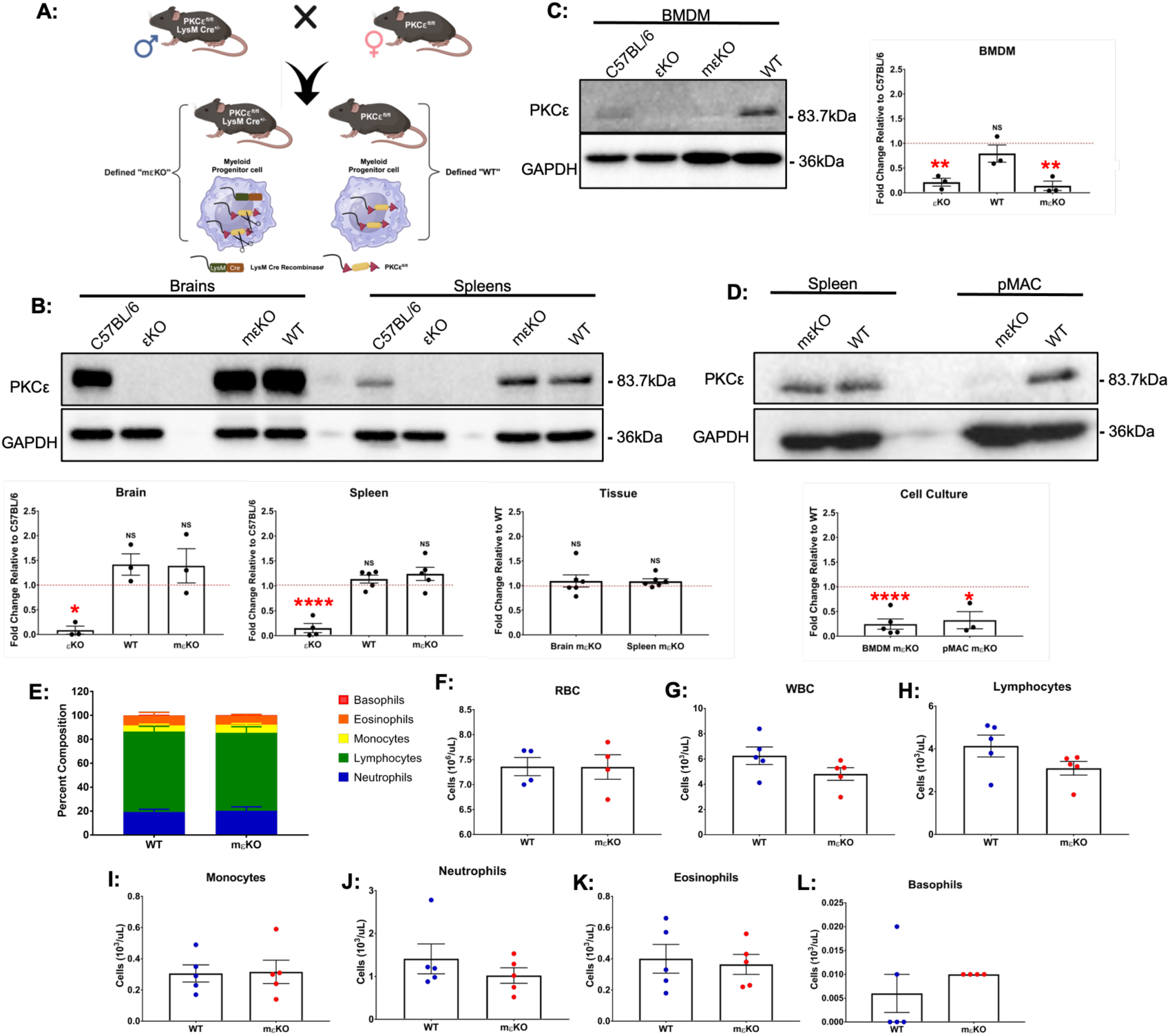
LysM Cre deletes PKCε selectively from myeloid cells and does not affect blood composition. (**A**) Schematic of the LysM Cre PKCε^fl/fl^ breeding strategy. (**B-D**) Representative Western blot and corresponding densitometry of PKCε and GAPDH expression in brains and spleens (**B**), BMDMs (**C**), and pMACs (**D**) (n=3-6). Tissues from the global PKCε knockout mouse (εKO) are included for reference; ratios of expression. No substantive reduction in PKCε expression was detected in brains or spleens of LysMCre expressers; little/no PKCε was detected in lysates from BMDMs or pMACs expressing Cre. (**E-L**) Blood was collected and analyzed with a Heska HT5 CBC analyzer. (**E**) Composition of total white blood cells (WBC). Total numbers of (**F**) Red blood cells (RBC), (**G**) WBC, (**H**) Lymphocytes, (**I**) Monocytes, (**J**) Neutrophils, (**K**) Eosinophils, and (**L**) Basophils are shown. Data are presented as mean ± SEM. Statistical analysis was performed by two-tailed unpaired t-test (**B,D,F-L**) and one-way ANOVA (**B,C,E**) Red dotted line indicates respective C57BL/6 or WT (ie, PKCε^fl/fl^) used as the normalization control. Each symbol represents an individual mouse. *\*p <* 0.05; *\*\*p <* 0.001; *\*\*\*\*p <* 0.0001.

To determine the efficiency of myeloid deletion, we compared PKCɛ protein expression across four mouse strains: C57BL/6 (parental, all cells positive), global PKCɛ knockout (ɛKO, all cells negative^54^; PKCɛ^fl/fl^ (designated WT, theoretically all cells positive), and LysM Cre^+/−^ PKCɛ^fl/fl^ (designated mɛKO: **<u>m</u>**yeloid selective PKC**<u>ɛ</u> <u>k</u>**nock**<u>o</u>**ut).^12^ Brains and spleens were used as controls; brain because PKCɛ expression is highest in this tissue and spleens because the majority of cells are lymphoid so PKCε expression should not be affected by LysM Cre. As predicted, PKCɛ was expressed in brains and spleens of C57BL/6 mice but was undetectable in their εKO counterparts (Figure 3B). PKCɛ expression in WT and mɛKO brains and spleens was not significantly different from C57BL/6 (Figure 3B) nor was mɛKO expression significantly different from WT (i.e., PKCɛ^fl/fl^). These results confirm the myeloid selectivity of LysM Cre deletion in the PKCε^fl/fl^ background (Figure 3B) and validate the specificity of the PKCε antibody.

With respect to the efficiency of PKCɛ deletion in MØ: BMDMs from mɛKO mice had no detectable PKCɛ; expression in mɛKO pMACs was reduced >85% compared to WT, a level not statistically different from mɛKO BMDMs (Figure 3C,D). In conclusion, PKCɛ expression in mɛKO brains and spleens is similar to the parental C57BL/6 tissues. In contrast, PKCɛ in mɛKO MØ is low/not expressed.

### Characterization of mεKO mice

Complete blood counts of WT and mεKO blood revealed no differences in total numbers of circulating red blood cells (RBC), white blood cells (WBC), lymphocytes, monocytes, neutrophils, eosinophils, or basophils (Figure 3E-L). These results parallel the more extensive immune cell analysis done on the global PKCɛ knockout mice (i.e., no differences in number or activity of any immune cell type.^54^ Additionally, we found no differences in the number of stem cells harvested from bones, the number of BMDM differentiated from WT and mεKO stem cells (p=0.76, n=6), or the number of elicited pMACs recovered from WT and mεKO (p=0.66, n=3), Finally, we found no differences in expression of FcγR, CD11b, MHCII and Ly6C between mɛKO and WT BMDM, suggesting that PKCɛ deletion does not alter MØ polarization (Supplemental Figures S1, S5D). These data suggest that, at steady state, deletion of myeloid PKCɛ has no substantive effect on the immune system.

To determine the effect of PKCε deletion on other PKCs, we compared their relative expression levels in BMDMs. Expression of PKC α, β δ, η, and ζ were not significantly different by PCR or western blotting (Figure 4 A-G). Thus, there is no obvious compensation for PKCɛ in mɛKO MØ. Additionally, the unique substrate specificity of PKCɛ.^15,51,54,55^ would make functional compensation difficult.

**Figure 4:**
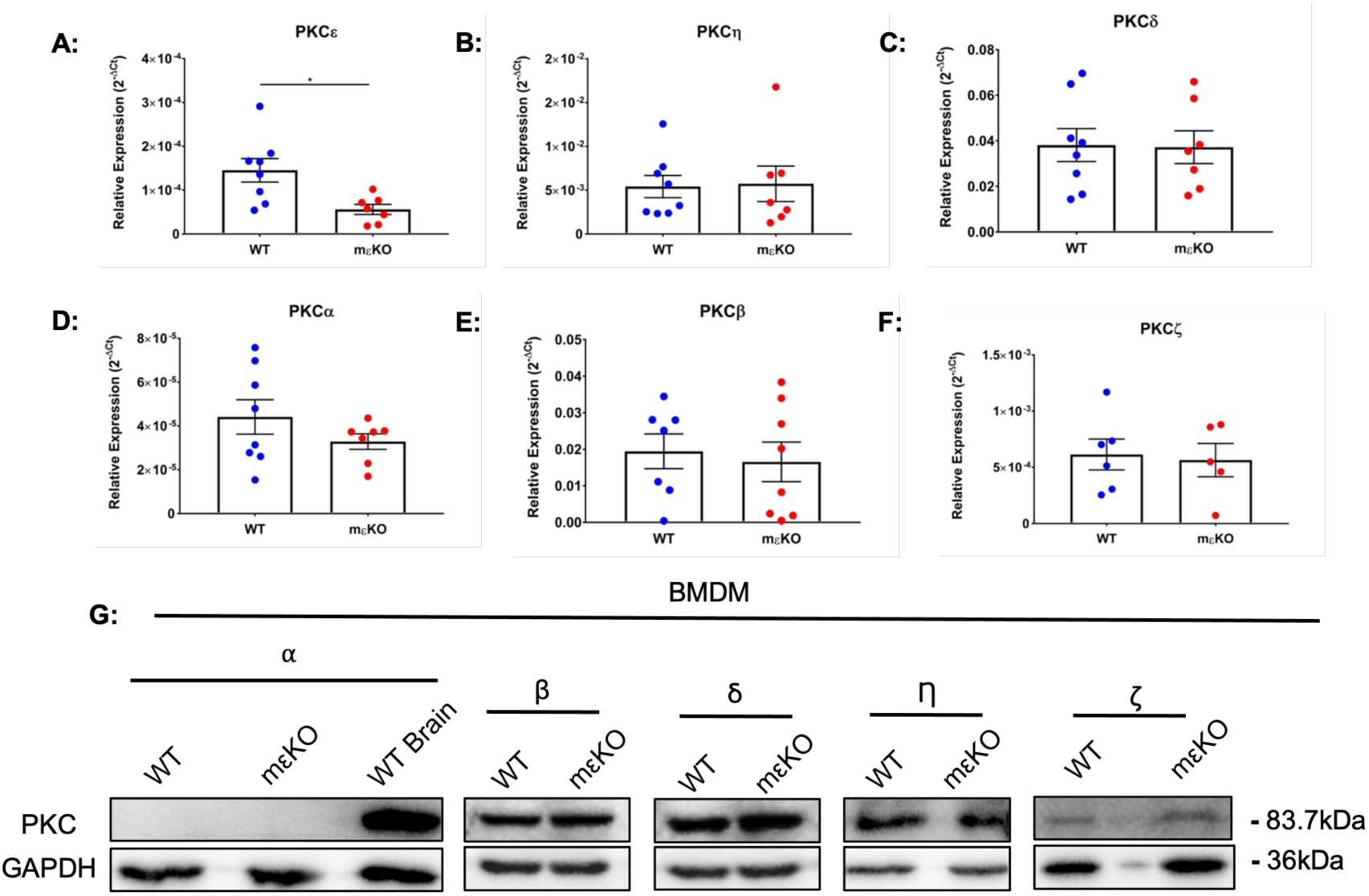
LysM Cre PKCε^fl/fl^ selectively deletes PKCε in MØ. (**A**) qPCR for novel (PKCε, PKCη, PKCδ), conventional (PKCα, PKCβ), and atypical (PKCζ) PKC’s reveal a significant decrease in PKCε with no effect on other isoforms, (n = 5-8, each symbol represents cells from a single animal). (**G**) Representative westerns confirming that PKCε deletion does not alter protein expression of other isoforms. (n=3). Data are presented as mean ± SEM. Statistical analysis was performed by two­tailed unpaired t-test in. *\*p <* 0.05.

### Deletion of PKCɛ does not affect expression of key atherosclerotic and inflammatory mediators

Since ɛKO BMDMs retained more and larger lipid droplets and released more TNFα (Figure 1), a likely explanation could be differential expression of scavenger receptors and/or a more polarized phenotype. qPCR of BMDM, elicited pMAC, and lipid loaded BMDM was used to determine the effect of PKCɛ deletion on the expression of 25 genes linked to atherosclerosis.^56^ Surprisingly, there were no statistical differences in message levels for any of the markers tested (Figure 5, Supplementary Figure S2-6). Similary, surface expression of scavenger receptors (SR-A1, SR-E2, SR-B2, SR-E1, SR-E2, and SR-B1) and pattern recognition receptors TLR 2 and 4 did not change with genotype or upon lipid loading (Figure 5, Supplementary Figure S2). Thus, any differences in plaque phenotype is more likely due to MØ PKCε signaling than intrinsic effects of differential receptor expression.

**Figure 5:**
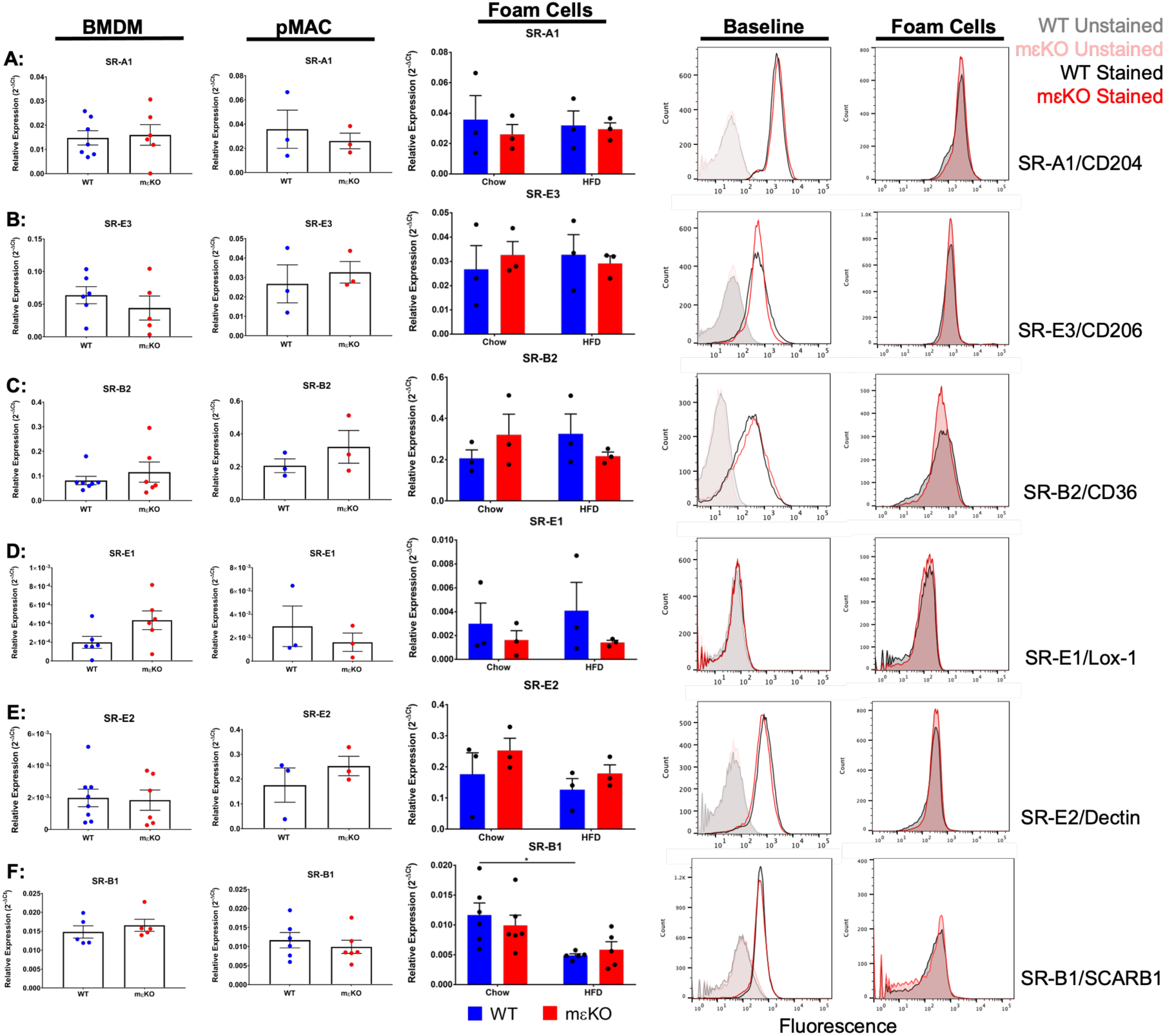
Deletion of PKCε does not alter scavenger receptor expression. (**A-F**) qPCR and flow cytometry for SR-AI (CD204), SR-E3 (CD206), SR-B2 (CD36), SR-E1 (Lox-1), SR-E2 (Dectin-1), and SR-B1 (SCARB1) in BMDMs (column 1,4) and elicited pMACs (column 2) at steady state (each symbol represents an individual animal, n = 3-6). In vivo derived foam cells (column 3, see Methods): qPCR of SR from elicited pMACs. In vitro derived foam cells (column 5): flow cytometry of SR receptors from BMDM loaded with Hi-oxLDL (50 µg/ml, 24 h). Flow cytometry are representative of 3-5 independent experiments. Data are presented as mean ± SEM. Statistical analysis was performed by two-tailed unpaired t-test and two-way ANOVA (there were no significance difference in expression between genotype).

### Plaques from hypercholesterolemic mɛKO mice are more advanced than WT

Knowing that MØ lacking PKCɛ retain more lipid (Figure 1A) and release more TNFα (Figure 1B), we hypothesized that loss of myeloid PKCɛ would exacerbate atherosclerosis *in vivo* (ie, PKCɛ expression is protective). AAV8-PCSK9 injections plus Western diet strategy^57^ was used to induce human-like hypercholesterolemia and plaques after 16 weeks (T=16, Figure 6A) Weight was recorded at the start of the experiment and prior to tissue harvesting (T =0 and 16 weeks). While weight increased with time, there were no significant genotype differences (Figure 6B). Likewise, plasma cholesterol (taken at weeks 2 and 16) increased with time but there were not genotype differences (Figure 6C); ≥500mg/dL of cholesterol was defined as hyper-cholesterolemic.^58^

**Figure 6:**
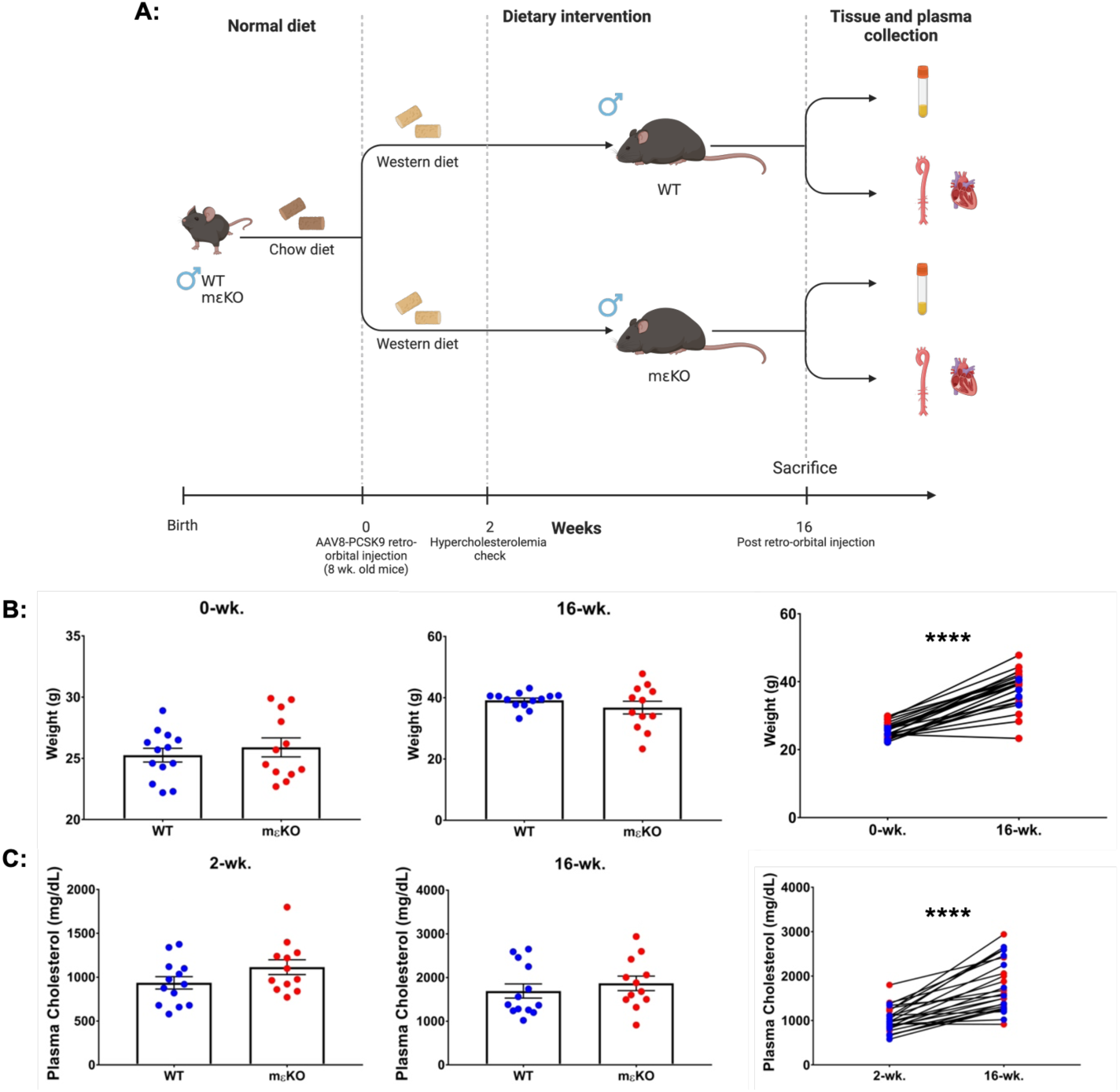
WT and mεKO animals have similar weights and plasma cholesterol levels in an in vivo model of hypercholesterolemia. (**A**) Timeline of in vivo model. Created in BioRender. Wells, A. (2024) https://BioRender.com/e98l871. (**B**) Weights were recorded at T = 0 and T = 16 wks. Genotype did not affect weight gain. (**C**) Blood was collected at 2 and 16 weeks for determination of plasma cholesterol concentrations. While cholesterol increased over the time course, there was no genotype difference. Each symbol (blue-WT, red-mεKO) represents one animal (n =12-13, two independent cohorts). Data are presented as mean ± SEM. Statistical analysis was performed by two-tailed paired t-tests (**B,C**) and unpaired t-tests (B,C). *\*\*\*\*p <* 0.0001.

Standard metrics of atherosclerosis include plaque area, foam cell content, amount of necrosis, and collagen cap thickness. Plaque area was measured in both the arotic root and aortic arch while quantitation of foam cell number, necrotic area and cap thickness was scored for arotic roots.

Lesion area in the aortic arch was quantified by Oil red O staining and normalized to arch area as detailed in Methods. Percent lesion coverage was significantly higher in mɛKO aortic arches compared to WT, indicative of a higher plaque burden (Figure 7A,C). To test for inter-observer variability, two observers quantified a subset of the aortic arches; no significant variability was found (Figure 7B).

**Figure 7:**
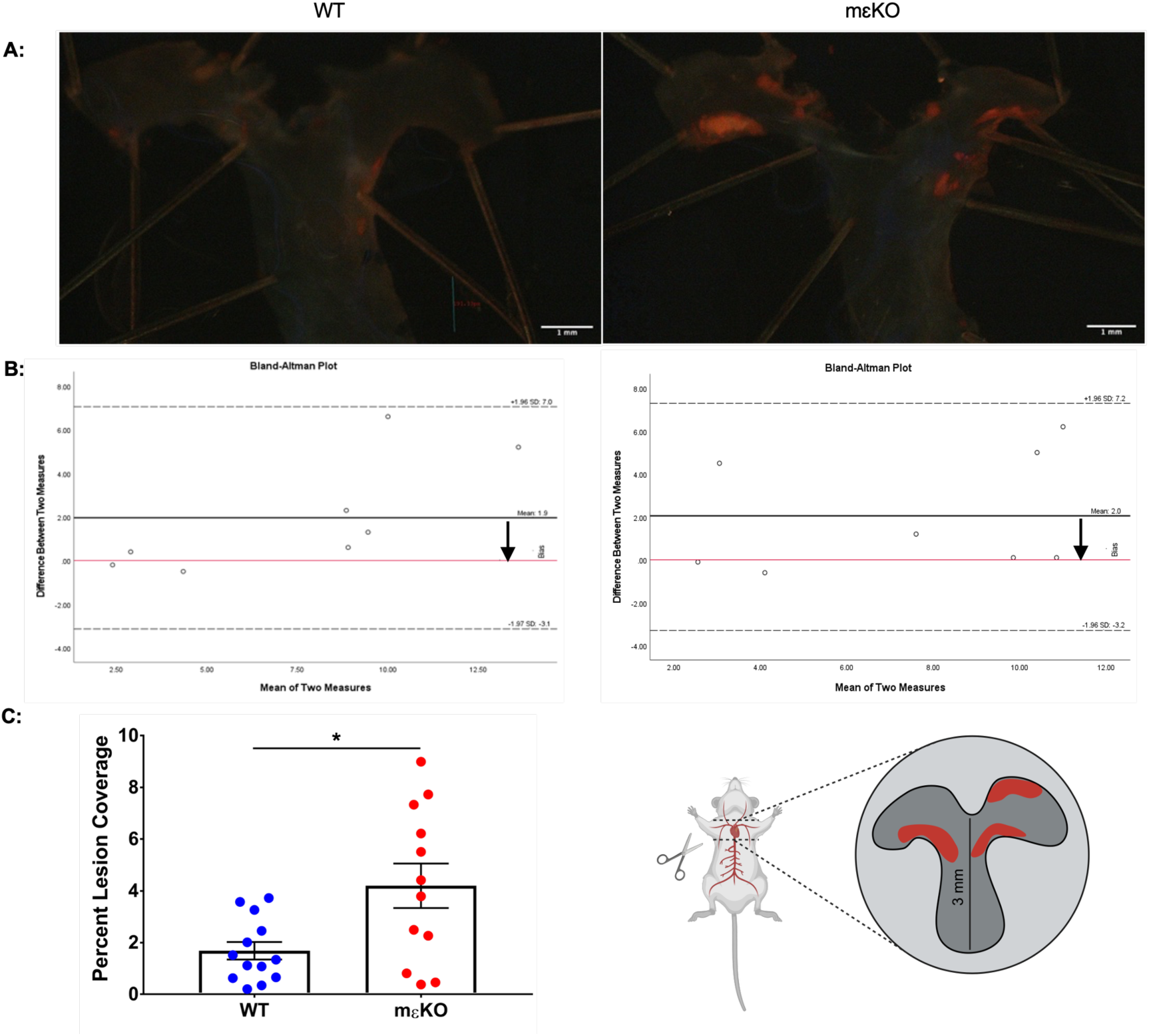
mεKO aortic arches have a higher percent lesion coverage than their WT counterparts. (**A**) Schematic of the region of descending aorta removed; area analyzed is enlarged. Created in BioRender. Wells, A. (2024) https://BioRender.com/q65k186. Representative images of Oil red O stained *en face* preparations of aortic arches recovered from animals after 16 weeks of hypercholesterolemia. (**B**) A subset of (8) mice were assessed for inter-observer variability, no shift in the bias was detected. (**C**) Quantitation of the aortic arch and 3 mm of descending aorta for percent lesion coverage (n = 12-13, each point represents one animal). See methods for details. Data are presented as mean ± SEM. Statistical analysis was performed by Bland-Altman analysis in (**B**) and two­tailed unpaired t-test in (**C**). Scale Bars: 1 mm. *\*p <* 0.05.

The aortic root lends itself to more detailed analyses. Sections were collected at the first sign of plaque near the aortic valve and sectioned such that each slide contained 3 × 10 µm sections at levels 250 µm apart (details in Methods). Six slides containing sections closest to the center of valves were stained with Hematoxylin and Eosin (H&E). Consecutive slides were stained for ORO, followed by Sirius red, followed by F4/80.

Lesion area and percent necrosis was quantified from H & E stained slides and recapitulated the results of the aortic arches in that the aortic root lesion area was significantly larger in mɛKO than WT. Necrotic area, as defined by H&E dull/devoid of nuclei, was significantly larger inmɛKO plaques (Figure 8A). Finally, we noted cholesterol clefts present in mɛKO plaques;few were apparent in WT plaques (Figure 8A). While difficult to quantify, embedded cholesterol crystals are a hallmark of advanced human plaques. ^59,60^

**Figure 8:**
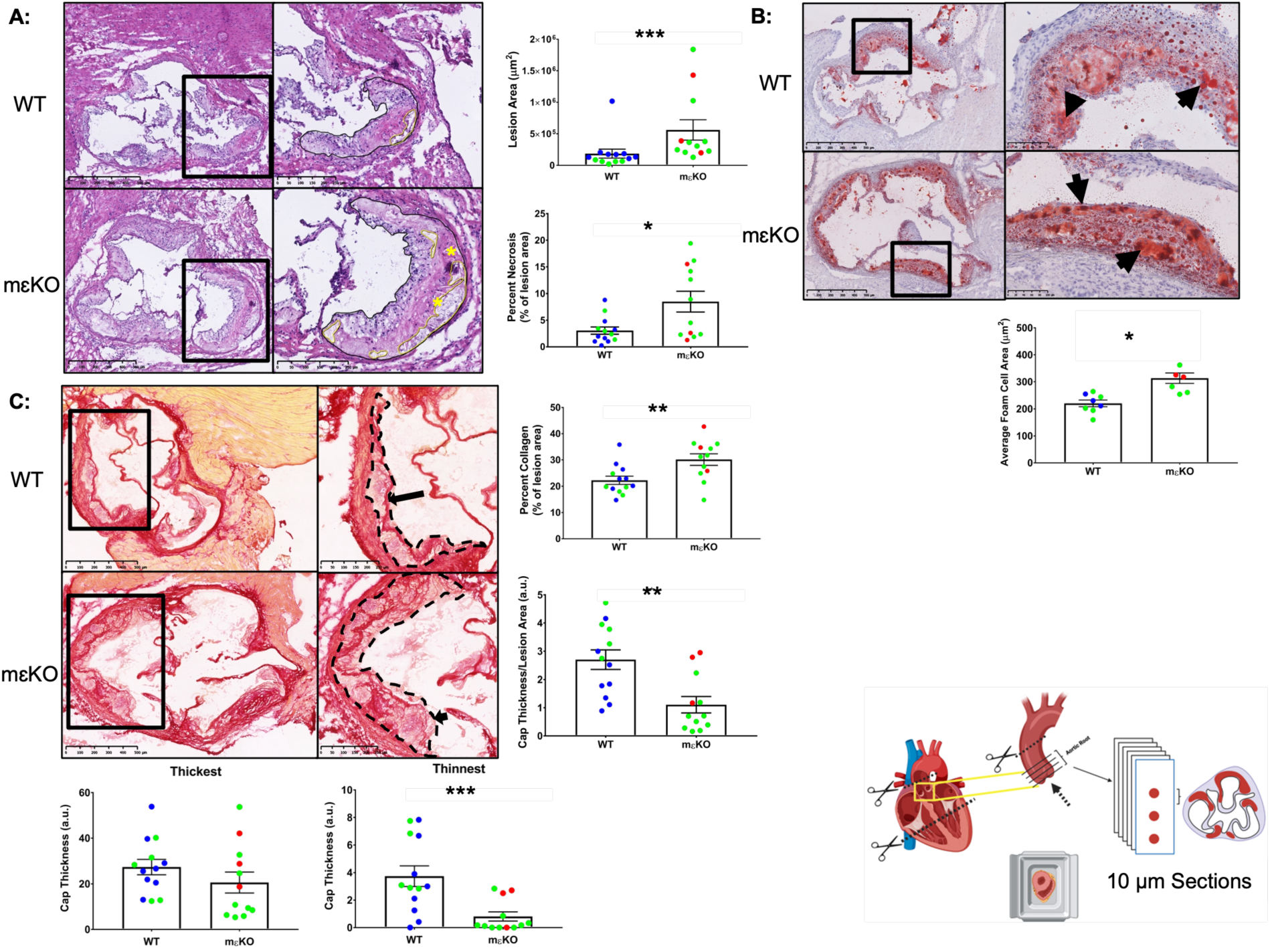
mεKO plaques have a more vulnerable phenotype. Animals were identified by ear tag such that the operator was blinded to genotype until all data was collected (n=12-13/genotype, 2 independent cohorts). See methods for processing/staining details. (**Bottom right**) Schematic of aortic root processing. Created in BioRender. Wells, A. (2024) https://BioRender.com/f19H90. (**A**) H&E staining: Plaques (black outlined), necrotic regions (yellow outlined) and cholesterol clefts (yellow asterisk) were identified and lesion area and percent necrosis per mouse were quantified. Plaques from mεKO mice were significantly larger and more necrotic. (**B**) Slides consecutive to H&E were stained with ORO and the average foam cell area per mouse was recorded; foam cells (black arrowheads) are significantly larger in mεKO plaques (n = 7-8). (**C**) Plaque sections were stained for collagen with Sirius Red, were outlined (black hashed lines) and overall collagen content reported as a percent of plaque area. As vulnerable plaques have thin collagen caps, we measured the average cap thickness per mouse as well as the thickest and thinnest regions; mεKO plaques have significantly thinner regions than their WT controls; arrow denotes a plaque with a cap and arrowhead denotes a plaque with no cap. (n = 12-13). (**A,C**) Scale bars: 500 µm (left) or 250 µm (right). (**B**) Scale bars: 500 µm (left) or 100 µm (right). Each symbol represents one animal and green symbols represent animals that are genotypically ZsGreen Positive. Data are presented as mean ± SEM. Statistical analysis was performed by two-tailed unpaired t-test in. *\*p <* 0.05; *\*\*p* < 0.01 ; *** p < 0.001.

Consecutive slides were stained with ORO to assess lipid retention. ORO rich areas were quantified, normalized to the number of nuclei (obtained from the contiguous H&E stained section), and reported as average foam cell area per mouse. Consistent with the larger plaque area, mεKO plaques have larger foam cells (Figure 8B). Larger foam cells would imply more retained lipid, providing in vivo validation of the in vitro lipid loading of PKCε negative MØ shown in Figure 1A. These data support the hypothesis that PKCɛ plays a role in lipid handling.

Consecutive slides were stained with Sirius red to visualize collagen. Overall content was quantified and normalized to lesion area. Surprisingly, there was significantly *more* collagen in mɛKO lesions compared to WT (Figure 8C). However, closer examination revealed that while collagen overlayed the foam cells in WT, much of it was buried in mɛKO samples. We previously reported buried collagen in carotid mouse plaques^43^, a metric that was used as one criteria to stage murine carotid plaques. “Buried” fibrosis is also seen in human plaques and is consistent with a previous disruption (i.e., instability).^61^ Thus, vulnerable plaques could have more collagen but with thinner caps. Larger and more necrotic plaques with little to no cap are indicators of vulnerability in humans; rupture is less likely if the soft necrotic core is protected by a thick fibrous cap. Applying this metric, we found significantly thicker collagen caps in WT plaques; many mɛKO plaques had regions with *no* cap (Figure 8C). Notably, a plaque is only as stable as its thinnest cap, therefore we assessed the thickest and thinnest plaque cap per mouse. There were and found no differences in the thickest cap regions. In contrast, mɛKO plaques have significantly thinner caps than their WT counterparts (Figure 8C).

Knowing MØ release MMPs that degrade the fibrous cap, and that MMPs 8, 9, and 12 are significantly upregulated in the inflamed regions of human plaques, we asked if MØ are the cells overlaying the collagen in mɛKO plaques.^62^ To do this, we exploited the availability of Jackson Laboratories **RCL-**ZsGreen reporter mouse and introduced this gene into some individuals in the mεKO colonly; cells with an active LysM Cre gene will be green. All the BMDM and pMACs from mɛKO mice expressed ZsGreen while their siblings lacking Cre did not (Figure S7B). Knowing that ZsGreen does not alter plaque metrics (Figure 8) and mɛKO MØ do not express PKCɛ (Figure 3C,D), we asked if the cells overlaying collagen were F4/80 positive (i.e., MØ). Plaques from ZsGreen^+/−^ mɛKO and WT were stained for F4/80. We found co-localization of ZsGreen and F4/80 in mɛKO but not WT mice (where Zs isn’t expressed), confirming that the cells overlaying collagen in mɛKO plaque are PKCɛ null MØ (Supplemental Figure S7A).

### Lipid loaded MØ from mɛKO mice retain more cholesterol

Histologically, mεKO plaques have larger foam cells than WT (Figure 8B). Also, we know that ɛKO BMDM retain more lipid than their C57BL/6 counterparts (Figure 1A). Given this information, we asked if mɛKO MØ have a lipid handling defect. To test this, WT and mɛKO BMDM were treated with highly-oxidized LDL (Hi-ox-LDL, 24 hours) and retained total (TC) and free cholesterol (FC) were quantified. Cholesterol ester (CE), the primary component of lipid droplets was calculated and reported as the difference between TC and FC. mɛKO BMDMs contained significantly more TC and CE (Figure 9A). These data utilize a different technique for quantifying lipid while providing an independent but consistent result that expression of PKCε moderates lipid retention by MØ. Indeed, these in vitro results are recapitulated in vivo (i.e, higher ORO staining in mεKO aortic roots; Figure 8B).

**Figure 9:**
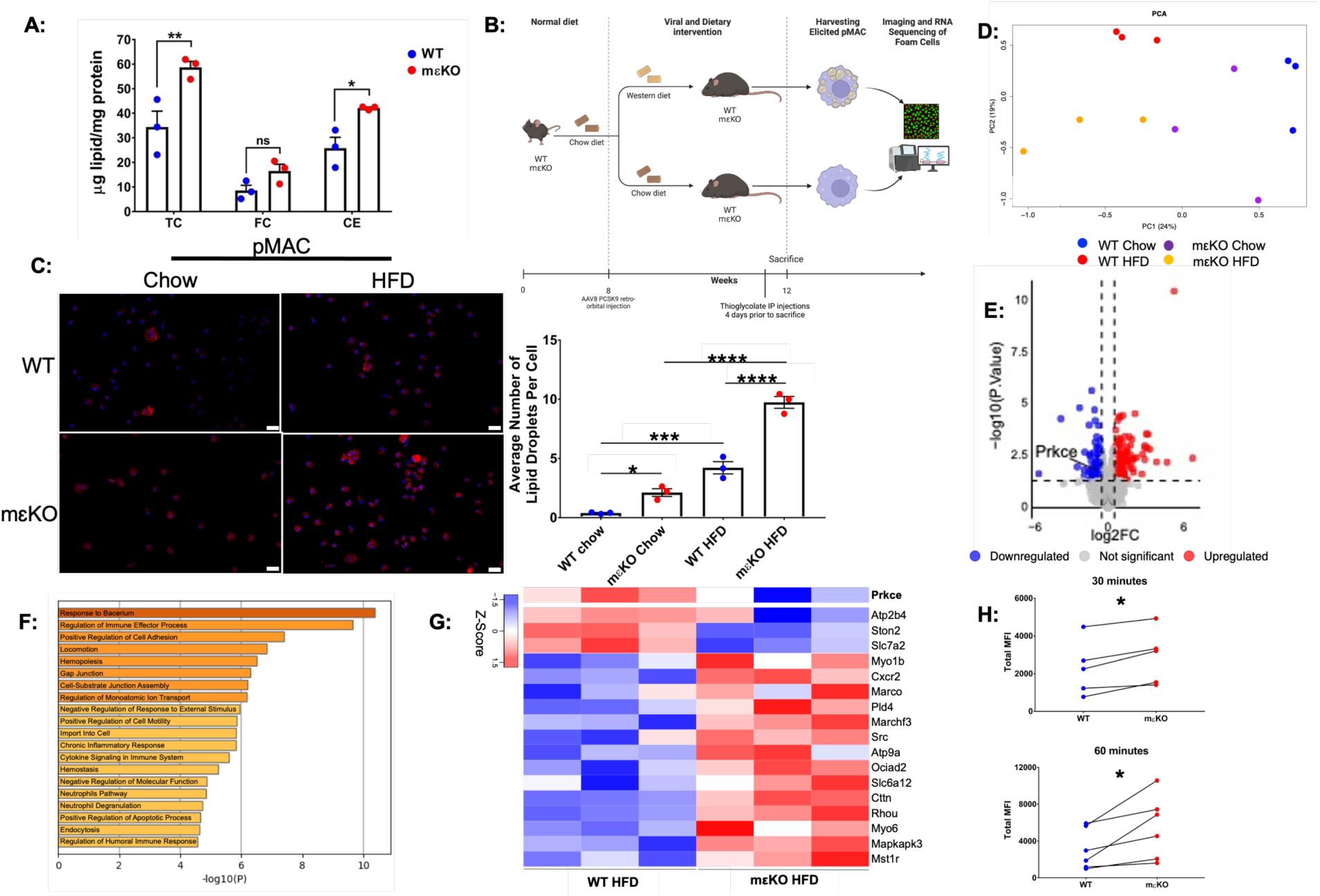
mεKO BMDMs and elicited pMACs retain more cholesterol/lipids in vitro and in vivo through enhanced endocytosis mechanisms. (**A**) BMDMs were loaded with human Hi-ox-LDL for 24h and assessed for total cholesterol (TC), free cholesterol (FC), and cholesterol esters (CE) (n = 3 with 4 technical replicates per n). (**B**) Strategy for generation of foam cells in. vivo. Created in BioRender. Wells, A. (2025) https://BioRender.com/f20m105. (**C**) Elicited pMACs from animals in (**B**) were stained with LipidTOX and Dapi to label neutral lipids and nuclei. Scale bar: 1 µm. The total number of well-defined lipid droplets was assessed by FIJI, normalized to the total number of nuclei, and reported as the average number of lipid droplets per cell. (**D-G**) RNA sequencing of elicited pMACs from hypercholesterolemic mice revealed significant differences in gene expression. (**D**) Principal component analysis (PCA) indicates distinct gene expression profiles as shown by differential clustering by genotype and diet. (**E**) Volcano plot from total RNA sequencing illustrating differentially regulated genes upon deletion of PKCε in hypercholesterolemic mice, where PRKCE is in relation. P value < 0.05 and fold change Iog2 > 2. (**F**) Gene Ontology analysis of differentially regulated genes upon PKCε deletion. (**G**) Heatmap of gene expression values for significantly downregulated and upregulated genes for import into the cell (i.e., endocytosis, phagocytosis, and receptor-mediated endocytosis) upon deletion of PKCε. (H) Synchronized endocytosis of Alexa-488 Ac-LDL for 30 and 60 minutes and assessed for the total MFI of lipid loaded (green) endosomes, (n = 3). Each point in **A-D,H** represents one animal n = 3-4. Data are presented as mean ± SEM. Statistical analysis was performed by one-way ANOVA (**A,C**) or paired t-test (H). *\*p <* 0.05; *\*\*p* < 0.01 ; *\*\*\*p* < 0.001, *\*\*\*\*p <* 0.0001.

While larger foam cells in mεKO plaques implies more lipid, it is not a direct measure of MØ lipid content. Thus we isolated elicited peritoneal macrophages from mεKO and WT mice that were injected with AAV8-PCSK9 and fed either a chow or high fat diet (4 weeks, Figure 9B). mεKO PMACs, even from animals on a chow diet, contained significantly more lipid droplets than WT (Figure 9C). While a HFD increased lipid retention in both genotypes, retention was substantially higher in mεKO cells (Figure 9C). In conclusion, these data provide in vivo validation of the in vitro results and implicate MØ PKCε in the regulation of regulates lipid dynamics in a way that could slow the progression of atherosclerosis. *<u>How</u>* this occurs is an open question.

### RNAseq

Given that mεKO MØ retain dramatically more lipid than WT, and that lipid loading does not alter SR (Figure 5) or efflux transporter expression (Supplemental Figure S3C,D), we are left with a conundrum: *How* is PKCε regulating lipid dynamics independently of receptor expression? We turned to RNAseq as a discovery based means to provides clues as to PKCε’s mechanism of action. We submitted the RNA from pMACs isolated from mɛKO and WT mice (Figure 9B) for RNA sequencing and produced the GSE291828 database. Principle Component Analysis (PCA) revealed distinct differences between chow and HFD groups confirming that diet changes gene expression. That there are differences between WT and mɛKO HFD animals indicates that genotype is also affecting gene expression (Figure 9D). To determine *how* PKCɛ may be protective in a hypercholesterolemic environment, we focused on the data from HFD fed mice. A volcano plot reveals 173 genes upregulated and 105 genes down-regulated in hypercholesterolemic mεKO compared to WT. PRKCE is significantly downregulated in mɛKO confirming the efficiency/selectivity of our LysM Cre PKCɛ^fl/fl^ model (Figure 9E). Not surprisingly, many of the upregulated gene families include response to bacteria and regulation of immune effector functions, consistent with the fact that the global PKCε knockout mouse succumbs to infections that are cleared by WT mice.^54^ Of particular interest is the Gene Ontology (GO) summary family *Import Into Cell* (Figure 9F). This family includes genes related to phagocytosis, endocytosis, and receptor-mediated endocytosis (Figure 9G), which dovetails with our previous work demonstrating a role for PKCɛ in phagocytosis, membrane mobilization, and vesicular trafficking.^12,13^ As modified lipids are taken up by endocytosis, and endocytosis would not necessarily require changes in receptor expression, we asked if *endocytosis* is altered in mεKO MØ. To test this, we implemented a model of synchronized endocytosis (30, 60 minutes) using Alexa-488 labeled Ac-LDL. At these early timepoints, mɛKO MØ contain significantly more fluorescent endosomes than their WT counterparts (Figure 9H) consistent with the hypothesis that PKCε regulates lipid trafficking independent of receptor expression. Further work is needed to elucidate exactly what that pathway is and how it is impacted by expression or absence of PKCɛ but this body of work is the first to implicate MØ PKCɛ in restricting/slowing atherosclerosis.

In summary, the overall composition of mɛKO plaques (larger, more necrotic, engorged foam cells, embedded collagen overlayed with new plaque, and significantly thinner caps) is consistent with human metrics of vulnerability. In addition, RNA sequencing reveal atheroprone mɛKO mice have increased expression in genes related to endocytosis and endocytosis studies confirm differences in the amount of endocytosis lipid (Figure 9H). As the mɛKO animals have PKCɛ deletion restricted to myeloid cells, we can conclude that loss of *myeloid* PKCɛ exacerbates the atherosclerotic program. Conversely, this work identifies myeloid PKCɛ as a potential atheroprotective gene.

## DISCUSSION

Knowing that FcR ligation stimulates TNFα release, and that FcR ligation activates PKCε, it was initially surprising that PKCε deletion enhanced TNFα release but had no effect on IL6 (Figure 1B). In contrast, ɛKO macrophages have blunted responses to LPS/IFNγ, including lower production of NO, TNFα, and IL6. ^54^ These conflicting results suggest that MØ responses are tuned to their environment, with their signaling response differing if initiated through TLR *vs* FcR. In the context of atherosclerosis, the scale maybe tipped in favor of an FcR activation pathway, based on the elevation of modified LDL-immune complexes in patients with atherosclerosis.^46,47,63^ Notably, Murray and Stow^64^ reported that secretion of TNFα and IL6 occurs by different pathways in MØ. Although the involvement of PKCɛ was not addressed, we would postulate that TNFα release involves PKCε signaling while IL6 does not. Furthermore, the fact that εKO secreted significantly more TNFα suggests PKCε expression moderates TNFα secretion, thereby limiting inflammation. According to GSE104140, PKCε expression is significantly decreased in advanced plaques and can be recapitulated in vitro (Figure 2B,C). Granted, this is a single database. Extensive searching of the GSE databases turned up one other curated database of human atherosclerotic plaques (GSE120521). The numbers of specimens were low (n=4 stable and 4 unstable). PKCε levels were not significantly different between stable vs unstable plaques. However, it should be noted that these were endarterectomy specimens; disease was already advanced to the point of intervention. With no control to compare, these specimens may reflect the gene signature of the advanced plaques of GSE104140. Indeed, PKCε is significantly lower in both advanced plaque stages of GSE104140 compared to diffuse intimal thickening (Fig 2B) but they are not significantly different from each other (although expression in advanced soft plaques trends lower). However, plaque tissue is a complex mix of cells. Thus, we tested PKCε knockout MØ for lipid uptake and cytokine release in response to FcR ligation (Fig 1). Deletion of PKCε exacerbates both of these metrics and led us to hypothesize that PKCε slows plaque progression.

Because PKCε is expressed in many tissues, and modulates such diverse cellular processes as endothelial function, smooth muscle cell migration, proliferation, and inflammation,^24,65,66^ determining its role in MØ required generation of a mouse lacking myeloid PKCε. LysM Cre was chosen as it is the most selective and efficient myeloid deletor;^53^ the specificity of LysM for PKCε deletion in myeloid cells paralleled that reported for other genes (Figure 3B-D, 4). We exploited the AAV8 PCSK9 + western diet strategy to mimic the hypercholesterolemia produced in humans carrying gain-of-function PCSK9 mutations^51,67^ rather than crossing into the more traditional ApoE or LDLR knockout mice.

We did note a light PKCε band in pMAC, but not BMDM lysates from mɛKO animals (Figure 3C,D) This signal may be from resident peritoneal B-1A cells that express high levels of PKCɛ (ImmGen Probe Set 10447294). Importantly, we established that, similar to the global PKCε knockout mouse, loss of MØ PKCε does not alter steady state blood composition (Figure 3E-L) or expression of 25 genes linked to atherosclerosis (Figure 5 and Supplemental Figures S2-S6). Thus, any differences in the context of hypercholesterolemia can be linked to myeloid PKCε.

Having established that plaques from mεKO mice are significantly more advanced than WT, and exhibit markers similar to that of vulnerable human plaques, we turned to lipid loaded MØ to probe mechanism (Figures 6-8). Lipid loading was done either in vivo (pMAC from hypercholesteremic mice) or in vitro (Hi-ox-LDL fed BMDM). Regardless of loading strategy, expression of the cohort of genes was unchanged, yet mεKO retained more TC and CE as well as more lipid droplets compared to control, arguing that PKCε regulates lipid handling but not through these genes (Figures 5, 9A-C).

RNAseq of pMAC isolated from hypercholesterolemic WT and mεKO mice provided a discovery-based approach to uncover potential signaling pathways impacted by PKCε expression. Consistent with results from the global knockout mouse, response to bacterium was dramatically upregulated in lipid loaded mεKO pMAC (Figure 9F). Potentially relevant to atherosclerosis are signaling pathways that would facilitate MØ entry into plaques (i.e., cell adhesion, cell motility, locomotion, chronic inflammation, and cytokine signaling). Additionally, upregulation of apoptosis signaling suggests that mεKO MØ may be more sensitive to apoptosis. This is in line with numerous cancer studies in which overexpressed PKCε was found to be anti-apoptotic.^32–37^ Notably, increased MØ apoptosis in plaques can lead to necrosis which is higher in mεKO plaques (Figure 8).

That loss of PKCε results in more lipid retention was validated in BMDM from mεKO and εKO mice and in pMACs isolated from hypercholesterolemic mice (Figures 1,9). This could theoretically be extended to humans as PKCε is lower in advanced plaques (Figure 2B). However, with no change in SR expression (Figure 5) and no apparent difference in expression of the efflux transporters ABCA1 and ABCG1 (Supplemental Figure S3C,D), we are left to consider PKCε signaling. We know that PKCε controls vesicular trafficking for membrane expansion during FcR mediated phagocytosis.^12,68^ Most lipid loading experiments rely on long term (4-72 hours) incubations for lipid droplet assessment and uptake typically occurs by receptor mediated endocytosis. To compliment this, RNA sequencing revealed the GO families *Import into Cell* and *Endocytosis* to be significantly upregulated in mεKO pMAC from hypercholesterolemic mice (Figure 9F,G). To test the hypothesis that endocytosis may be elevated in mεKO MØ, we followed uptake of A488 labeled Ac-LDL for 30 and 60 minutes. At these early timepoints mεKO MØ were significantly more fluorescent that their WT counterparts (Figure 9H), consistent with enhanced endocytosis.

Given no change in receptor expression, our data suggests PKCε acts on an intracellular signaling network; its loss leads to increased lipid retention. But, we know that PKCε regulates vesicle delivery from Golgi to phagosomes and that vesicles are not delivered in PKCε null MØ.^12,68^ How then, do we explain the apparent disparity, its loss decreases vesicle trafficking to the plasma membrane in phagocytosis but increases lipid uptake during hypercholesterolemia? One possibility is that PKCε functions at a trafficking node to move vesicles along microtubules. During phagocytosis, it brings Golgi-derived vesicles to the phagosome and phosphorylates proteins necessary for vesicle fusion.^12^ It may facilitate endocytosis by regulating trafficking and/or fusion along the maturation path. Its loss may interrupt endosome maturation, allowing accumulation of lipid laden endosomes and ultimately, larger foam cells. That mεKO MØ have more endosomes after 30 minutes of lipid loading (Figure 9H), retain more TC and CE (Figure 9A), and lipid droplets after lipid loading (Figure 9C), is consistent with such a model. While requiring a rigorous test of the hypothesis, this work provides a framework for studies going forward.

In conclusion, we generated and characterized the mεKO mouse with PKCε selectively and efficiently deleted in myeloid cells. Utilizing the AAV8-PCSK9 + western diet strategy, we established that mεKO mice have more advanced plaques with markers (e.g. thin/no collagen caps, more necrosis, cholesterol clefts) reminiscent of vulnerable human plaques. mεKO MØ retain more lipid despite no differences in expression of SR or ABCA1 ABCG1. RNAseq results highlight endocytosis as a signaling pathway upregulated in MØ from hypercholesterolemic animals and endocytosis is significantly elevated in mεKO MØ. Together, these results are the first to report a protective role for PKCε in atherogenesis. The reduction of PKCε in advanced human plaques suggests that strategies to retain PKCε expression may represent a novel therapeutic approach to limit atherosclerosis and the medical conditions associated with cardiovascular disease. Our ongoing challenge is to determine the mechanism of PKCε protection and why it decreases as atherosclerosis progresses.

## Supporting information

Wells et al Supplemental

## Non-standard Abbreviations and Acronyms

CVD: Cardiovascular Disease
MØ: Macrophages
LDL: Low Density Lipoprotein
Hi-ox-LDL: Highly Oxidized LDL
Ac-LDL: Acetylated LDL
LD: Lipid Droplets
MI: Myocardial Infarction
BMDM: Bone marrow derived macrophages
pMAC: Peritoneal Macrophages
PKCε: Protein Kinase C epsilon
C57BL/6: C57BL/6 Mouse Strain
εKO: Global PKCε Knockout Mouse Strain
WT: PKCε^fl/fl^ Mouse Strain
mεKO: LysM Cre PKCε^fl/fl^ Mouse Strain
AAV8-PCSK9: Recombinant Adeno-Associated Viral Vector Serotype 8 – Proprotein Convertase Subtilisin/Kexin Type 9
HFD: High Fat Diet/Western diet
OCT: Optimal Cutting Temperature Compound
ORO: Oil Red O
TNFα: Tumor necrosis factor-α
IL6: Interleukin 6
TC: Total Cholesterol
FC: Free Cholesterol
CE: Cholesterol Esters
EDTA: Ethylenediaminetetraacetic Acid
DMEM: Dulbecco’s Modified Eagle’s Medium
BMM: Bone Marrow Media
FBS: Fetal bovine serum
PFA: Paraformaldehyde
PBS: Phosphate-buffered saline
ANOVA: Analysis of Variance
SEM: Standard error of the mean
PCA: Principle Component Analysis
CBC: Complete Blood Count

## Acknowledgments

The authors thank Dr. Gabrielle Fredman for her expertise in atherosclerosis and constructive feedback on experimental design, technical training, and data analysis. Dr. Younghwa Goo for help with cholesterol assays. Thank you to Dr. Paul Feustel for aiding in statistical analysis. The authors would also like to thank Genome Editing and Animal Models Core at the University of Wisconsin (Madison) Biotechnology Center for supplying the PKCɛ^fl/fl^ sperm, Taconic Biosciences (Rensselaer, NY) for producing PKCɛ^fl/fl^ mice, Cheryl Zjad for doing the initial animal husbandry and aiding in the development of the LysMCre PKCɛ^fl/fl^ murine line, Rosemary Prestipino for assisting in the creation of the graphical abstract, Albany Medical College’s Bioinformatic Core for producing graphs from RNA sequencing data, and BioRender for aiding in diagram development. ATW backcrossed the PKCɛ^fl/fl^ mice to generate the congenic strain for Jackson Labs (B6(Cg)-*Prkce^tm1.1Akv^*/LennMmjax stock# 69556), maintained the LysM Cre PKCɛ^fl/fl^ colony, designed and conducted experiments, analyzed the data, and wrote the manuscript. RBR managed and analyzed NCBI’s geneset GSE104140 and our GSE291828. MMS processed and analyzed the aortic arch tissue and ran qPCR and western blots. RHB processed MØ and ran qPCR. AED ran and analyzed experiments using the global (ɛKO) knockout MØ. MRL coordinated the research, provided the funding, ran and analyzed flow cytometry, and edited the manuscript.

## Sources of Funding

This work was supported by R03 AI144841, R21 AI185827, Johnathan Vasiliou Foundation, and Albany Medical College Bridge Funds.

## Disclosures

None.

## Supplemental Material

Figure S1-S7

Full unedited gels

Major Resources Table

## Notes

The authors have declared that no conflict of interest exists.

### Competing Interest Statement

The authors have declared no competing interest.

### Summary of Updates

Addition of RNAseq data and information on mechanism.

