## Supplementary material for "Identification of Myeloid Protein Kinase C – Epsilon as a Novel Atheroprotective Gene": Wells et al Supplemental

**Supplemental Figure Count:** 7
**Major Resources Table**

**Methods**

**F4/80 staining and analysis.**

Consecutive sections to those stained with Sirius red were taken to assess if MØ in mεKO plaques lack PKCε. Four sections spaced 40 μm intervals were dried at RT for 10 minutes, rehydrated in 1X PBS for 10 minutes, then fixed (4% PFA, 10 min). Roots were then blocked for non-specific staining (0.5% fish gelatin, 2% BSA, 0.3% Triton X-100 in 1X PBS) for 1 hour at RT. Tissue was probed with F4/80 monoclonal antibody, APC (Invitrogen, MA5-16625), diluted in blocking buffer according to manufacturer's instructions, overnight at 2-8°C. Slides were washed 3 times for 15 minutes each in 1X PBS, mounted with ProLong Glass Antifade Mountant with NucBlue (Invitrogen, P36981) and left to cure overnight prior to imaging. Merge and fire LUT images were created on FIJI software.

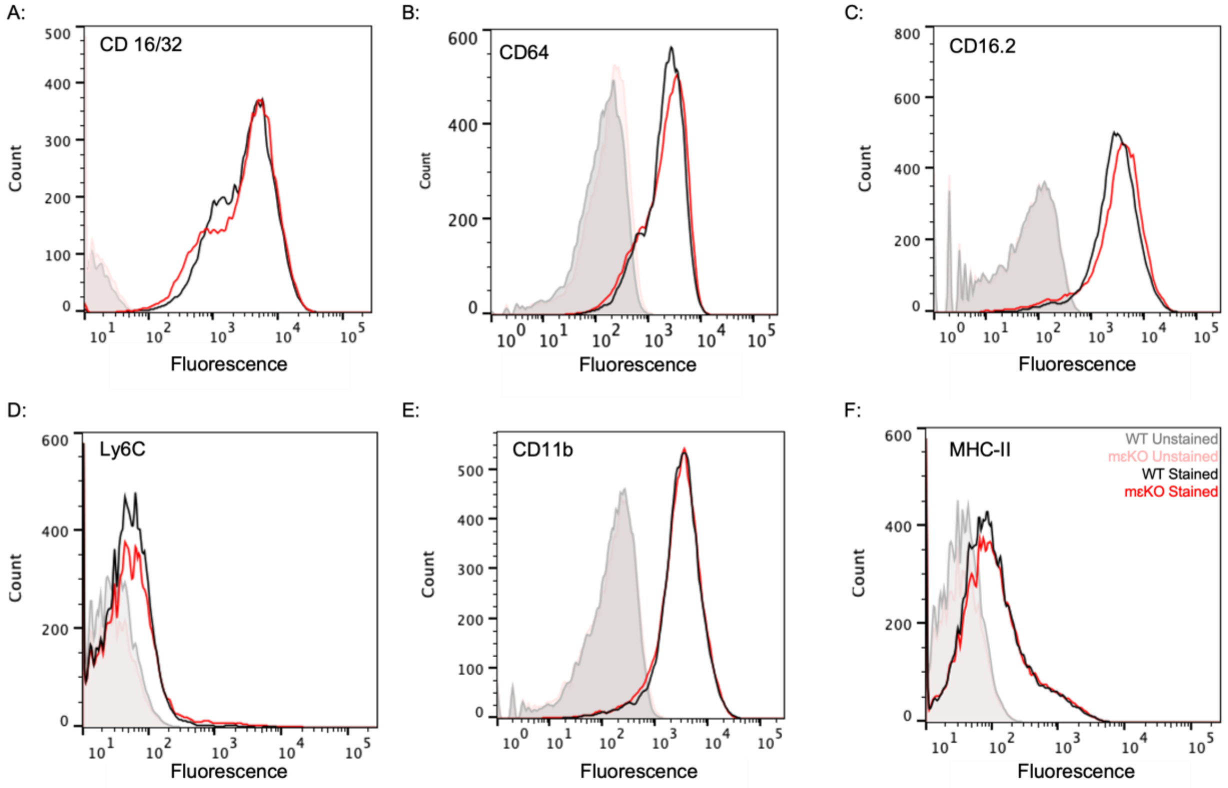


**Supplemental Figure S1: Myeloid PKCε deletion does not alter steady state expression of FcγR, CD11b, or MHC-II.** Representative flow cytometry plots for expression of surface CD16/32 - FCγRII/III (**A**), CD64 – FCγRI (**B**), CD16.2 – FCγRIV (**C**), Ly6C (**D**), CD11b (**E**), and MHC-II (**F**) in WT (black/gray) and mεKO (red/pink) BMDMs (n = 3). Methods detailed in manuscript.

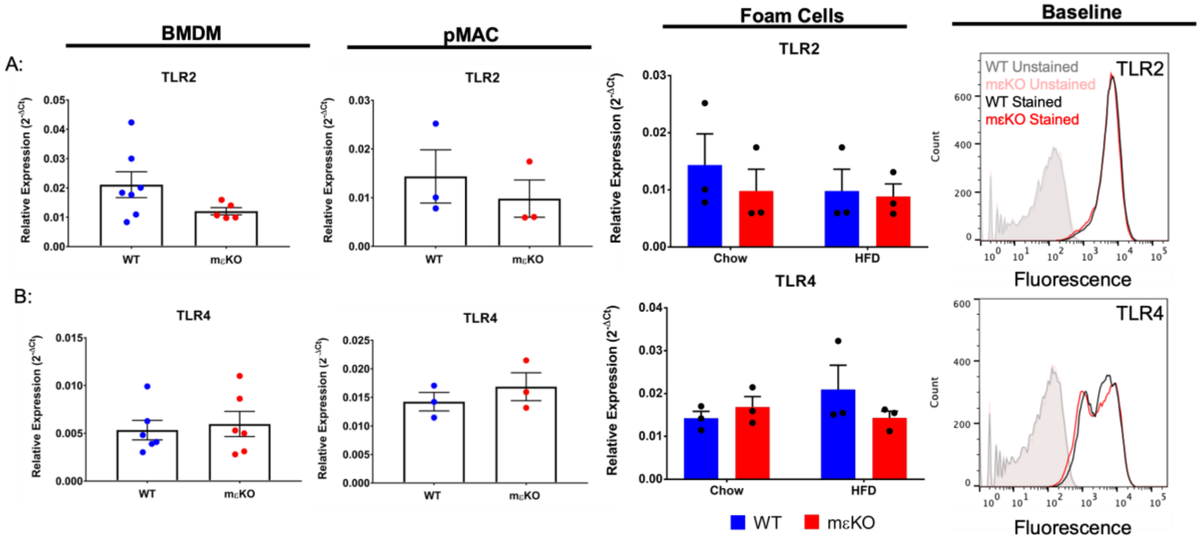


**Supplemental Figure S2: Deletion of PKCε does not alter expression of TLR2 or TLR4.** (**A,B**) qPCR for TLR2 and TLR4 in BMDMs (Column 1), elicited pMACs (Column 2), and elicited pMAC foam cells – in vivo loaded (Column 3) (n = 3-6). Flow cytometry for surface receptor expression (n = 2). Data are presented as mean ± SEM. Statistical analysis was performed by two-tailed unpaired t-test. Methods detailed in manuscript.

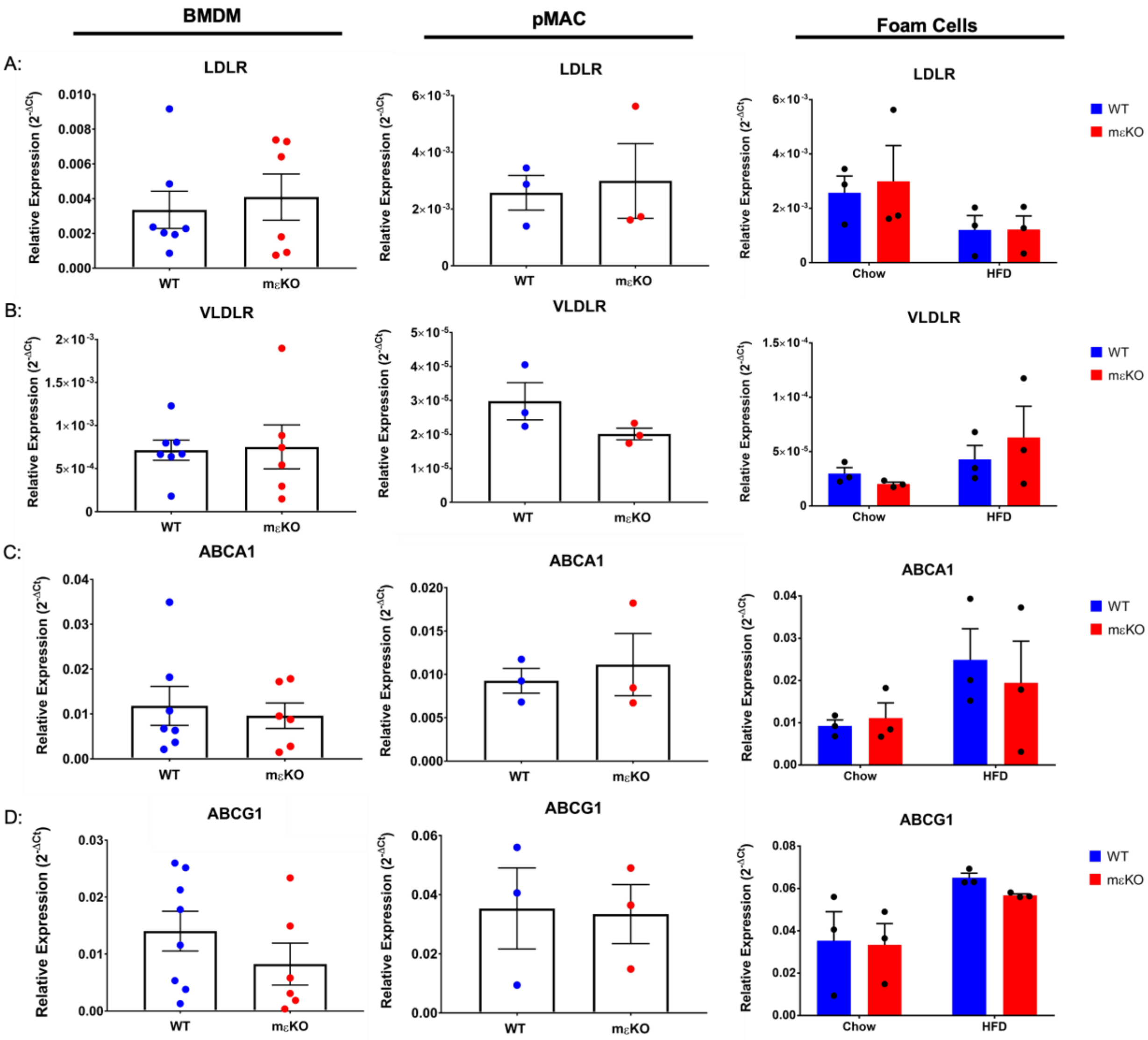


**Supplemental Figure S3: Deletion of PKCε does not alter expression of lipid uptake/efflux receptors.** (**A-D**) qPCR for LDLR, VLDLR, ABCA1, and ABCG1 in BMDMs (Column 1), elicited pMACs (Column 2), and elicited pMAC foam cells – in vivo loaded (Column 3) (n = 3-6). Data are presented as mean ± SEM. Statistical analysis was performed by two-tailed unpaired t-test. Methods detailed in manuscript.

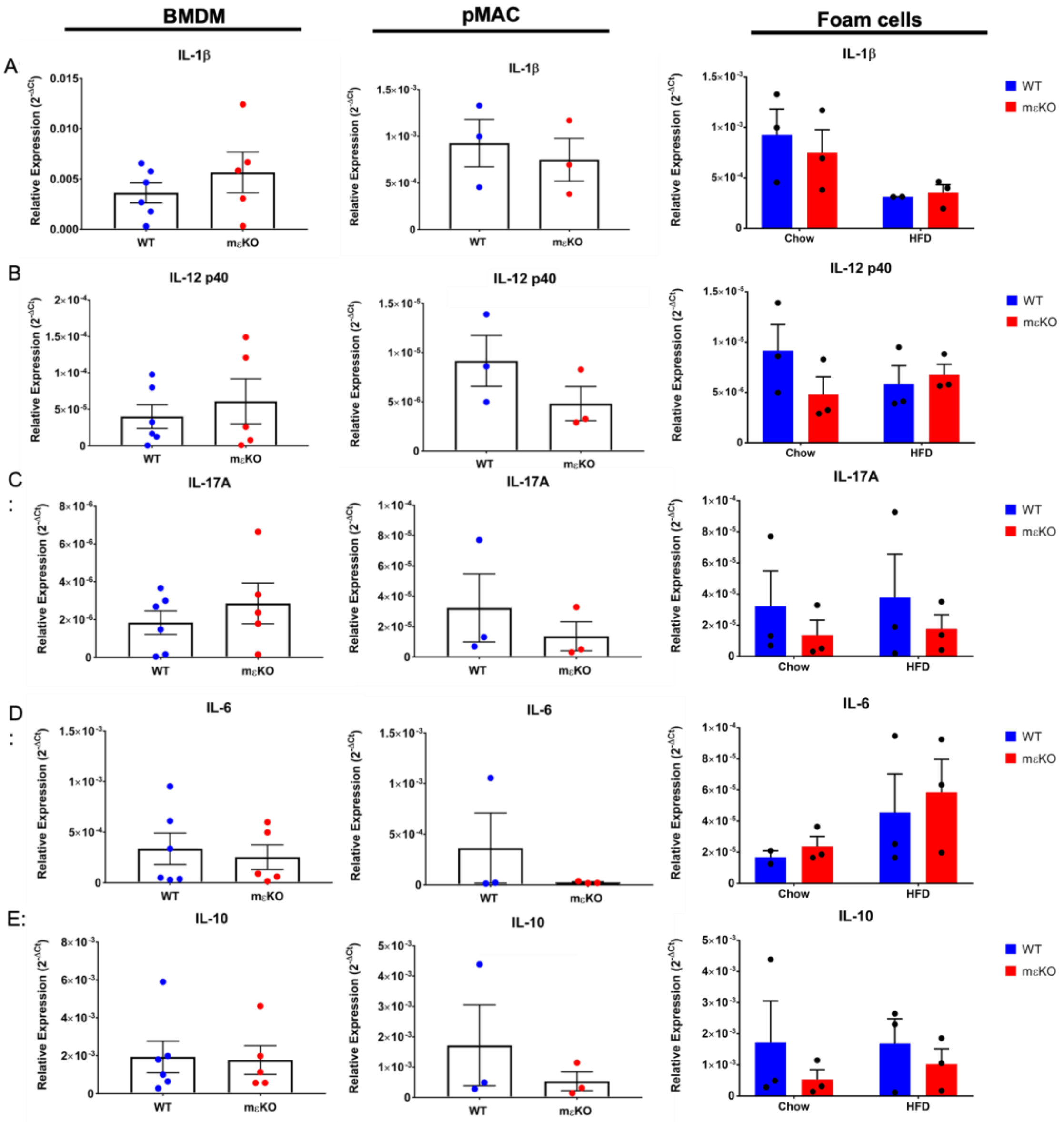


**Supplemental Figure S4: Deletion of PKCε does not alter cytokine expression.** (**A-E**) qPCR for IL-1β, IL-12 p40, IL-17A, IL-6, and IL-10 in BMDMs (Column 1), elicited pMACs (Column 2), and elicited pMAC foam cells – in vivo loaded (Column 3) (n = 3-6). Data are presented as mean ± SEM. Statistical analysis was performed by two-tailed unpaired t-test. Methods detailed in manuscript.

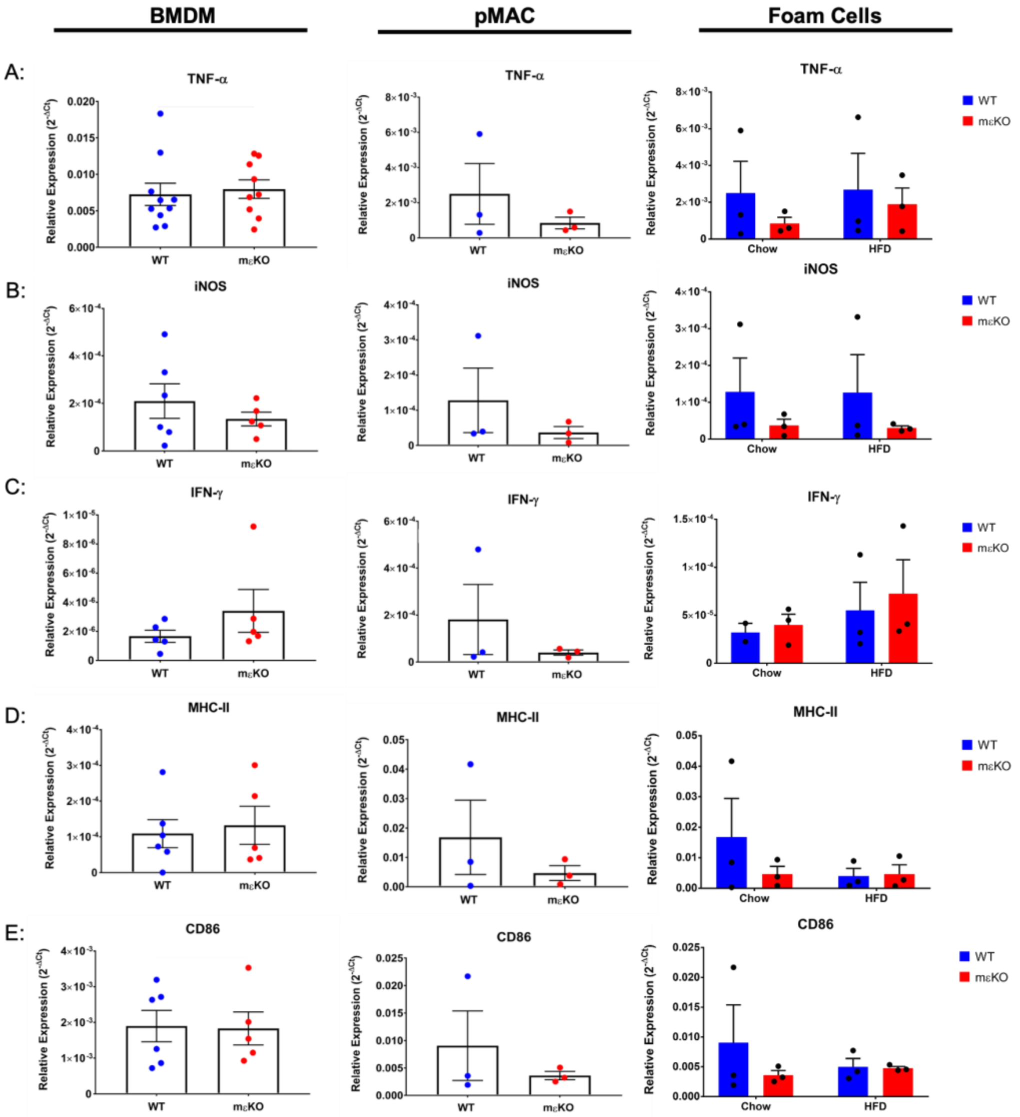


**Supplemental Figure S5: Deletion of PKCε does not alter expression of M1 polarization markers. (A-E)** qPCR for TNFα, iNOS, IFN-γ, MHC-II, and CD86 in BMDMs (Column 1), elicited pMACs (Column 2), and elicited pMAC foam cells – in vivo loaded (Column 3) (n = 3-6). Data are presented as mean ± SEM. Statistical analysis was performed by two-tailed unpaired t-test. Methods detailed in manuscript.

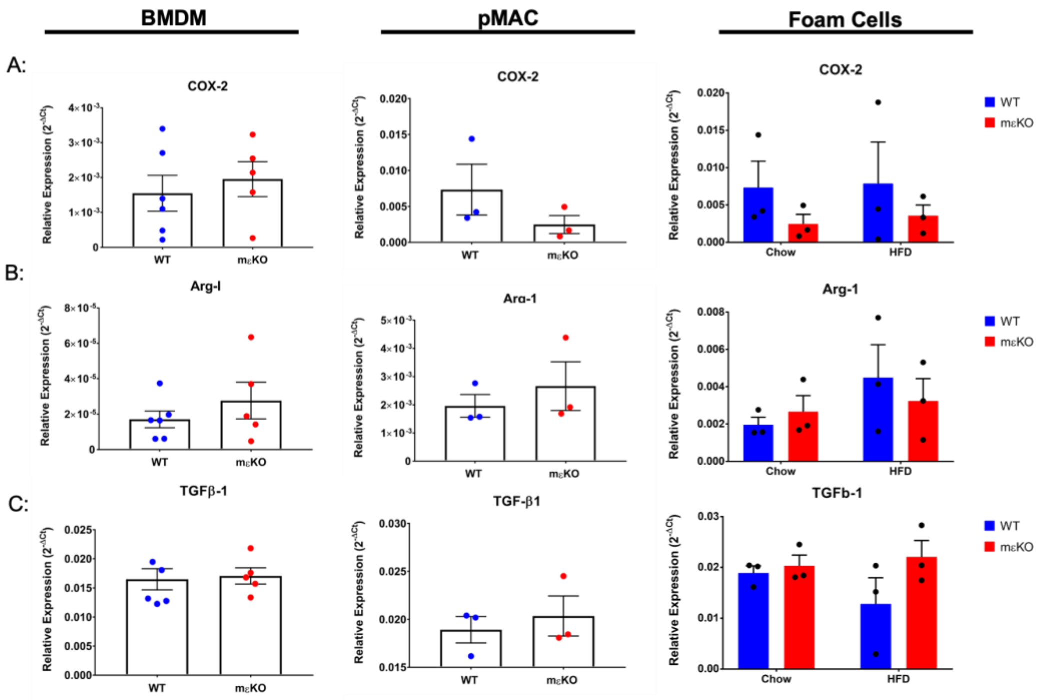


**Supplemental Figure S6: Deletion of PKCε does not alter expression of M2 polarization markers.** (**A-C**) qPCR for COX-2, Arg-1, and TGFβ-1 in BMDMs (Column 1), elicited pMACs (Column 2), and elicited pMAC foam cells – in vivo loaded (Column 3) (n = 3-6). Data are presented as mean ± SEM. Statistical analysis was performed by two-tailed unpaired t-test. Methods detailed in manuscript.

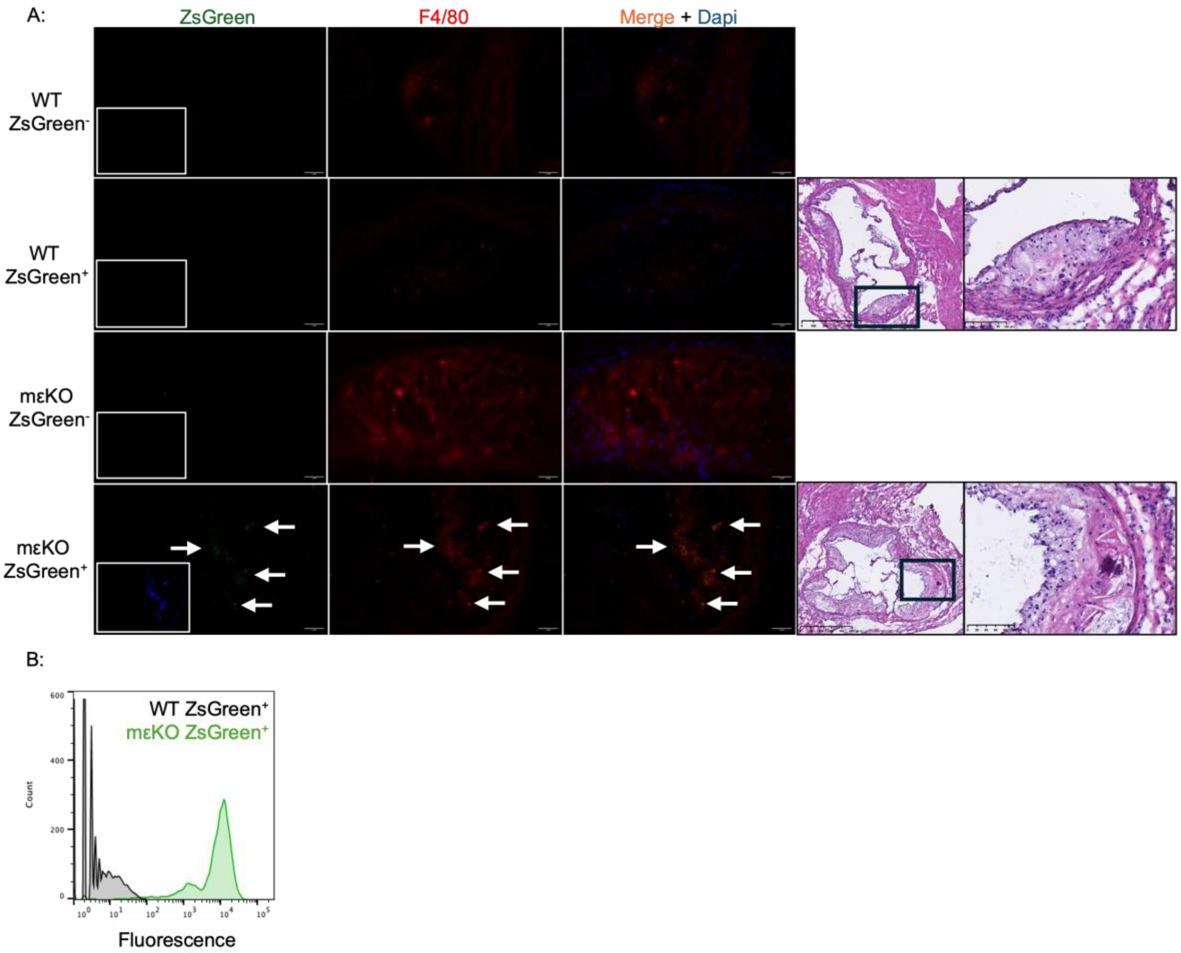


**Supplemental Figure S7: ZsGreen Cre reporter gene confirms Cre expressing MØ are in mεKO plaques, and they overlay mεKO plaques as new growth.** (**A**) Some of the colony carry the ZsGreen reporter gene, allowing detection of myeloid cells. Plaque from ZsGreen positive and negative animals were stained with F4/80 and dapi. Arrowheads indicate colocalized ZsGreen and F4/80. Green cells are detected in ZsGreen$^{+}$ mεKO, but not WT, animals. Insets are fire LUTs of ZsGreen. Sequential sections of ZsGreen$^{+}$ plaques were stained with H&E confirming the majority of cells present are MØ and are collected at the outer edge of mεKO plaques aas new growth but are encapsulated by a thick cap in WT. Representative of 3 animals per group. (**B**) Flow cytometry of BMDM from WT and mεKO animals carrying the ZsGreen reporter confirm that mεKO, but not WT, MØ express the reporter. Representative of 3 independent experiments. (**A**) Scale bars: 2 µm for immunohistochemistry; 500 µm for H&E whole root; 100 µm for magnified region of plaque used for F4/80 staining.

### Major Resources Table

#### Genetically Modified Animals

| Name | Species | Vendor or Source | Background Strain | Name/Stock Number |
|---|---|---|---|---|
| **C57BL/6** | Mouse | Jackson | C57BL/6 | C57BL/6 stock# 000664 |
| **εKO** | Mouse | Jackson | C57BL/6 | *B6.129S4-Prkce$^{tm1Msg}$/J* stock# 004189 |
| **WT** | Mouse | Jackson | C57/BL6 | B6(Cg)-*Prkce*$^{tm1.1Akv}$/LennMmjax stock# 69556 |
| **mεKO** | Mouse | Jackson | C57BL/6 | WT crossed with the Jackson Laboratory, Strain #004781 B6.129P2-*Lyz2*$^{tm1(cre)Ifo}$/J |

#### Western blot Antibodies

| Name | Type | Vendor | Catalog # | Dilution Factor |
|---|---|---|---|---|
| PKC-epsilon | Rabbit pAb | Millipore-Sigma | #06-991 | 1:500 |
| PKC-epsilon | Rabbit mAb | ThermoFisher Scientific | #MA5-14908 | 1:500 |
| PKC-delta | Rabbit mAb | Cell Signaling Technology | D10E2 #9616 | 1:2,000 |
| PKC-beta | Rabbit mAb | Cell Signaling Technology | D3E70 #46809 | 1:500 |
| PKC-alpha | Rabbit PC | Cell Signaling Technology | #2056 | 1:500 |
| PKC-eta | Rabbit mAb | AbCam | #ab179524 | 1:500 |
| PKC-zeta | Rabbit mAb | Cell Signaling Technology | C24E6 #9368 | 1:500 |
| GAPDH | Rabbit mAb | Cell Signaling Technology | #2118 | 1:10,000 |
| GAPDH | Mouse mAb | GeneTex | CTx627408 | 1:15,000 |
| Secondary | Goat anti-rabbit HRP (highly cross-adsorbed) | ThermoFisher Scientific | #A16110 | 1:10,000 |
| Secondary | Goat anti-mouse HRP (highly cross-adsorbed) | Invitrogen | #A16078 | 1:10,000 |

#### Flow Cytometry Antibodies

| Name | Fluorochrome | Vendor | Clone | Dilution (μl/5x10$^5$Cells) |
|---|---|---|---|---|
| CD11b | SBV515 | Bio-Rad #MCA711SBV515 | 5C6 | 2 |
| CD16.2 | Biotin | BioLegend #149512 | 9.00E+09 | 2 |

| | | | | |
|---|---|---|---|---|
| CD16/32 | BV421 | BD Biosciences #562896 | 2.4g2 | 1 |
| CD204/SR-A1 | APC | BioLegend #102610 | HM36 | 2.5 |
| CD206/SR-E3 | Biotin | LifeTech #MA5-16869 | MR5D3 | 2 |
| CD36/SR-B2 | A700 | Bio-Rad #MCA2748A700 | MF3 | 2.5 |
| CD64 | A647 | R & D Systems #FAB20741R | 290322 | 1 |
| Dectin-1/SR-E2 | PE | Bio-Rad #MCA2289PE | 2A11 | 2.5 |
| Lox-1/SR-E1 | Alexa 405 | R & D Systems #FAB1564V | 21402 | 2.5 |
| SCARB1/SR-B1 | Alexa 647 | NOVUS #NB400-104AF647 | Polyclonal | 4 μg/mL |
| Ly6C | SBV515 | Bio-Rad #MCA2389SBV515 | ER-MP20 | 2 |
| MHCII | SB780 | LifeTech #78-5321-82 | M5/114.15.2 | 0.05 |
| Streptavidin | SBV790 | Bio-Rad #STAR210SBV790 | Ø | 1 |
| TLR2 | PE | LifeTech #12-9021-80 | 6C2 | 0.05 |
| TLR4 | PE-Cy7 | BioLegend #145407 | SA15-21 | 1 |

#### RT-qPCR Primer Pairs

| Name | Sequence |
|---|---|
| β-actin | TCGCCTGAGGCTCTTTTCC<br>AGTTTCATGGATGCCACAGGAT |
| PKCε | GGGGTGTCATAGGAAAACAGG<br>GACGCTGAACCGTTGGGAG |
| PKCη | TCCGGCACGATGAAGTTCAAT<br>TACGCTCACCGTCAGGTAGG |
| PKCδ | ACATTCTGCGGCACTCCTGACT<br>CCGATGAGCATTTCGTACAGGAG |
| PKCα | ACAACCTGGACAGAGTGAAACTC<br>CTTGATGGCGTACAGTTCCTCC |
| PKCβ | CCAAGATGACGATGTGGAGTGC<br>CTCCATCACAAAGTACAGGCGG |
| PKCζ | GGCAGAGAAAACTTCCAGAGGAG<br>ATGTGTCCGTCGGCATCAAGGA |
| CD36/SR-B2 | GCCTCCTTTCCACCTTTTGTT<br>CGTAGATAGACCTGCAAATGTCAGA |
| Lox-1/SR-E1 | GGTTCCCTGCTGCTATGACTCT<br>GGCGTAATTGTGTCCACTGTACA |
| CD204/SR-AI | CAGACTGAAGGACTGGGAACACT<br>GGAGGCCCTTGAATGAAGGT |

| | |
|---|---|
| SCARB1/SR-B1 | ACACCCGAATCCTCGCTGGAAT<br>CCGTTGGCAAACAGAGTATCGG |
| CD206/SR-E3 | GGATGTTGATGGCTACTGGAGAA<br>GGGATTTCGTCTGATTTTTTGC |
| Dectin-1/SR-E2 | ACAATGCTGGCAACTGGGCTCT<br>AGAGCCATGGTACCTCAGTCTG |
| LDLR | CCAGACCCAGAGCCATCGTA<br>CGGGTGTTCCCCAATCTGT |
| VLDLR | CATCCTTCCTCTCTTGCTCTTAGTG<br>GCCAATTCCTCCACATCAAGTAG |
| TLR2 | CCCATTGAGAGGAAAGCCATT<br>CTCCAGGTAGGTCTTGGTGTTCA |
| TLR4 | CTTTATTCAGAGCCGTTGGTGTATC<br>CCAGAGCGGCTGCTCAGA |
| ABCA1 | GGGTGGTGTTCTTCCTCATTACTG<br>TTTACAGGTCTGGGCCTGATG |
| ABCG1 | GACACCGATGTGAACCCGTTTC<br>GCATGATGCTGAGGAAGGTCCT |
| iNOS | TGGAGCGAGTTGTGGATTGTC<br>CCAGTAGCTGCCGCTCTCAT |
| TNF-α | CATCTTCTCAAAATTCGAGTGACAA<br>TGGGAGTAGACAAGGTACAACCC |
| IL-12 p40 | CATCAGGGACATCATCAAACCA<br>TGACCTCCACCTGTGAGTTCTTC |
| IFN-γ | GGATGCATTCATGAGTATTGC<br>CCTTTTCCGCTTCCTGAGG |
| CD86 | TGTTCTGGAAACGGAGTCAATG<br>GGAGATGGAAACTCTTGAGTGAAATT |
| IL-6 | GCCAGAGTCCTTCAGAGAG<br>GGTCCTTAGCCACTCCTTC |
| IL-17A | GGACTCTCCACCGCAATGAA<br>GCACTGAGCTTCCCAGATCAC |
| COX-2 | TCATTCACCAGACAGATTGCTGG<br>TCCAAGCTCTACCATGGTCTCC |
| IL-1β | AGATGAACAACAAAAAAGCC<br>TCTATCTTGTTGAAGACAAACC |
| Arg-I | GTCTGGCAGTTGGAAGCATCA<br>TGTGAGCATCCACCCAAATG |
| TGFβ-1 | TGAACCAAGGAGACGGAATACA<br>GGAGTTTGTTATCTTTGCTGTCACAA |
| IL-10 | GATTTTAATAAGCTCCAAGACCAAGGT<br>TTCTATGCAGTTGATGAAGATGTCAA |
| MHCII | TCTGTCCTGGTGGCTCTG<br>TGGACGCATCAGCAAGGG |

#### Commercial Assays

| Description | Vendor/Catalog# | URL |
|---|---|---|

| | | |
|---|---|---|
| ELISA MAX Standard Set Mouse IL-6 | Biolegend #431301 | https://www.biolegend.com/en-us/products/mouse-il-6-elisa-max-standard-2250 |
| ELISA MAX Standard Set Mouse TNF-α | Biolegend #430901 | https://www.biolegend.com/en-us/products/mouse-tnf-alpha-elisa-max-standard-2242 |
| Cholesterol/Cholesterol Ester-Glo Assay | Promega Corporation #J3190 | https://www.promega.com/products/energy-metabolism/lipid-metabolism-assay/cholesterol-cholesterol-ester-glo-assay/?catNum=J3190 |

**Data & Code Availability**

| Description | Source | URL |
|---|---|---|
| iDEP | BMC Bioinformatics | https://bmcbioinformatics.biomedcentral.com/articles/10.1186/s12859-018-2486-6 |
| Trim Galore | GitHub | https://github.com/FelixKrueger/TrimGalore |
| Finctional enrichment analysis | WebGestalt | https://www.webgestalt.org/ |

**Statistical Software**

| Description | URL |
|---|---|
| GraphPad Prism | https://www.graphpad.com/features |
| SPSS | https://www.ibm.com/spss |
| R | https://cran.r-project.org/mirrors.html |

**Study Design**

| Groups | Sex | Age | Number (prior to experiment) | Number (after termination) | Littermates | Other description |
|---|---|---|---|---|---|---|
| Group 1 (WT) | M | 8 wks | 13 | 13 | Partial | Total number of animals are from two independent experiments. |
| Group 2 (mεKO) | M | 8 wks | 12 | 12 | Partial | Total number of animals are from two independent experiments. |

**Sample Size:** At least an n=4 for each experiment was required. More were included if they were the appropriate sex, genotype, and age matched (within the same week of birth).

**Exclusion Criteria:** Any mice with wounds, as well as runts were excluded from animal studies. In addition, females were excluded due to lack of sustained hypercholesterolemia. Future studies will titrate AAV8-PCSK9 doses.

**Randomization/Blinding:** All animals were chosen and processed in a random and blinded fashion. Animals were organized based on ear tag number. The code was broken once all raw data had been collected and was necessary to know genotype for statistical analysis.
